# Coherent multi-level network oscillations create neural filters to favor quiescence over navigation in *Drosophila*

**DOI:** 10.1101/2022.03.11.483976

**Authors:** Davide Raccuglia, Raquel Suárez-Grimalt, Laura Krumm, Cedric B Brodersen, Anatoli Ender, Sridhar R. Jagannathan, York Winter, Genevieve Yvon-Durocher, Richard Kempter, Jörg RP Geiger, David Owald

## Abstract

For all animals, undisturbed periods of rest are essential for undergoing recuperative processes. How neural interactions create brain states capable of dissociating an animal from its external world to promote quiescence remains a fundamental question. Here, we show how coherent network oscillations can create neural filters that favor a quiescent brain state over a state that promotes navigation. Circadian regulation and excitability of the *Drosophila* sleep homeostat (dFSB) generate nighttime specific slow-wave coherence between neural networks mediating sleep need (R5) and networks gating locomotion (helicon cells). Optogenetically mimicking coherent activity reveals that temporally fine-tuned R5 oscillations promote a quiescent state and reduce responsiveness to visual stimuli by hierarchically overruling locomotion-promoting helicon cells. We uncover that R5 and helicon bidirectionally regulate behavioral responsiveness by providing antagonistic inputs to head direction targets (EPG). Thus, coherent oscillations can form the mechanistic basis of neural filters by temporally associating antagonistic inputs and therefore reducing the functional connectivity between locomotion gating and navigational networks.

## Introduction

Whether we execute cue-driven behavior or remain unresponsive depends on the neural integration of internal states and the salience of sensory stimuli. External information is thought to be processed through efficient neural filtering mechanisms, which largely differ during wake and sleep, thereby defining gateways to the whole brain. Filters regulate the neural representation of sensory information and therefore create a switch that determines whether we enter an ‘allocentric’ mode directed to external stimuli or remain in an ‘egocentric’, internal-centered setting (Lu et al., 2022; Lyu et al., 2022). In mammals the thalamus has been implicated as a filter for sensory information, relaying it to higher order cortical networks (Andrillon and Kouider, 2020; Crunelli and Hughes, 2010; Gent et al., 2018). The lowest degree of behavioral responsiveness occurs during deep sleep, during which thalamo-cortical interactions generate slow-wave activity (SWA) that result from electrical coherence across cortical networks (Buzsaki and Draguhn, 2004). It is therefore conceivable that oscillatory activity patterns could be suited for providing a certain ‘cut-off’ while retaining the flexibility for transduction of salient or strong stimuli. Experimental evidence and knowledge on the mechanistic underpinnings however remain limited. Strikingly, SWA associated with sleep and sleep need has been observed in mammals (Siclari and Tononi, 2017), birds (Low et al., 2008), reptiles (Shein-Idelson et al., 2016), fish (Leung et al., 2019), crabs (Ramon et al., 2004) and recently in the vinegar fly *Drosophila melanogaster* (Raccuglia et al., 2019), and could represent an evolutionarily optimized mechanism for constructing neural filters. Indeed, mounting evidence suggests that neural coherence generating SWA is a network-autonomous marker of sleep need which might represent a fundamental neural principle for suppressing sensory processing to promote the reorganization of synaptic connectivity (Krueger et al., 2019; Suárez-Grimalt and Raccuglia, 2021; Vyazovskiy et al., 2011). However, how oscillatory coherence maintains quiescent states, gates signals to the brain and regulates behavioral responsiveness on a mechanistic network level remains largely unclear.

In *Drosophila*, behavioral responsiveness is particularly reduced during quiescent states, which are defined by prolonged inactivity (≥ 5 min) (Shaw et al., 2000). These quiescent states meet the criteria for sleep as they underlie homeostatic regulation (Shaw et al., 2000). Interestingly, transitions from an active state to a quiescent state are accompanied by an increase of oscillatory activity in the central brain (Yap et al., 2017). Recent years have established that sensory processing, behavioral responsiveness and sleep in *Drosophila* are integrated by a higher sensory processing center called the central complex (Donlea, 2019).

Importantly, in *Drosophila*, the individual circuit elements of the central complex are known and therefore allow a targeted analysis of network activity and its mechanistic relationship to quiescent states. While the network comprising the fan-shaped body (dFSB) functions as a homeostatic sleep switch (Donlea et al., 2009; Kempf et al., 2019; Pimentel et al., 2016), ExR1/helicon cells are part of a network which processes visual information and gates locomotion (Donlea et al., 2018). The R5 network, based on its connectivity likely engages in both navigation and sleep regulation (Blum et al., 2021; Flores-Valle et al., 2021; Huang et al., 2020; Liu et al., 2016; Raccuglia et al., 2019; Vafidis et al., 2022). Indeed, we recently discovered that in the R5 circuit, network-specific synchronization generating compound SWA arises over the day to integrate circadian and homeostatic sleep drive, promoting consolidated sleep phases to ensure adequate sleep quality (Raccuglia et al., 2019).

In this study, we directly target circuit elements of the central complex to investigate how the functional components of reciprocally connected networks (dFSB-helicon-R5) interact to create a filtering mechanism that maintains quiescence. Using a strategy that combines genetically-encoded voltage and Ca^2+^ sensors with optogenetics, we discovered that within this circuitry circadian and homeostatic sleep regulation are integrated to generate a brain state that is characterized by coherent SWA. Particularly, the locomotor-promoting helicon and sleep need mediating R5 circuit elements synchronize their electrical patterns during the night. Optogenetically mimicking these activity patterns during the day induced a quiescent state with reduced behavioral responsiveness to light. We further show that head direction neurons (EPG), which are part of a navigational system, receive antagonistic input from helicon and R5 to bidirectionally regulate behavioral responsiveness. We therefore propose that oscillatory network coherence provides the mechanistic basis for a neural filter by rhythmically associating antagonistic inputs, effectively reducing the functional connectivity between locomotion gating networks and the navigational system.

## Results

### Electrical slow-wave activity in the master sleep homeostat in *Drosophila*

In humans, sleep depth and mounting tiredness are associated with SWA (0.5-4 Hz) spreading across several brain regions (Adamantidis et al., 2019). We recently identified that the emergence of SWA in the R5 network of *Drosophila* is correlated to undisrupted sleep and reduced behavioral responsiveness (Raccuglia et al., 2019). Similar to thalamo-cortical circuits, the current *in vivo* brain state and the ability to generate SWA is maintained in the intact explant brain (Liu et al., 2016; Raccuglia et al., 2019; Sanchez-Vives and McCormick, 2000), enabling us to study activity patterns and network interactions characteristic for specific brain states. We therefore used this approach to investigate whether sleep-relevant SWA was a local phenomenon constrained to R5 neurons or also becomes manifest within other networks.

We first focused on the dFSB network which comprises neurons that integrate and execute sleep drive (Donlea et al., 2014). To directly test whether compound electrical activity is correlated between the dFSB and R5 network we performed dual color voltage imaging experiments by simultaneously expressing the green voltage indicator (GEVI) ArcLight (Cao et al., 2013; Raccuglia et al., 2016) and the red voltage indicator Varnam (Kannan et al., 2018) in R5 and dFSB, respectively (**Fig. 1A-C**). We focused on presynaptic compound activity because it represents the neural activity of the entire networks. We found that the presynaptic compound activity of R5 and the dFSB were largely synchronized at night (**Fig. 1D**). To verify interactions between these structures, we investigated whether diurnal variations of the electrical patterns observed in R5 were also observable in the dFSB. Indeed, compared to the morning we observed an increase in slow-wave power (0.5-1.5 Hz) at night when measuring the presynaptic compound signal, which represents the coherent activity of the whole dFSB network (**Fig. 1E, F**). Independent multi-cellular recordings revealed low overall activity of dFSB neurons in the morning hours (**Fig. S1A**). General activity markedly increased in the evening (**Fig. 1G**) and power analysis revealed an activity peak between 0.5-1 Hz (**Fig. 1H**). However, unlike what we previously observed for the R5 network (Raccuglia et al., 2019), activity was only loosely correlated between the single units regardless of whether measured during the morning or the night hours (**Fig. S1B**), indicating that changes in single-cell excitability are sufficient to drive diurnal changes at the network level.

**Figure 1.**
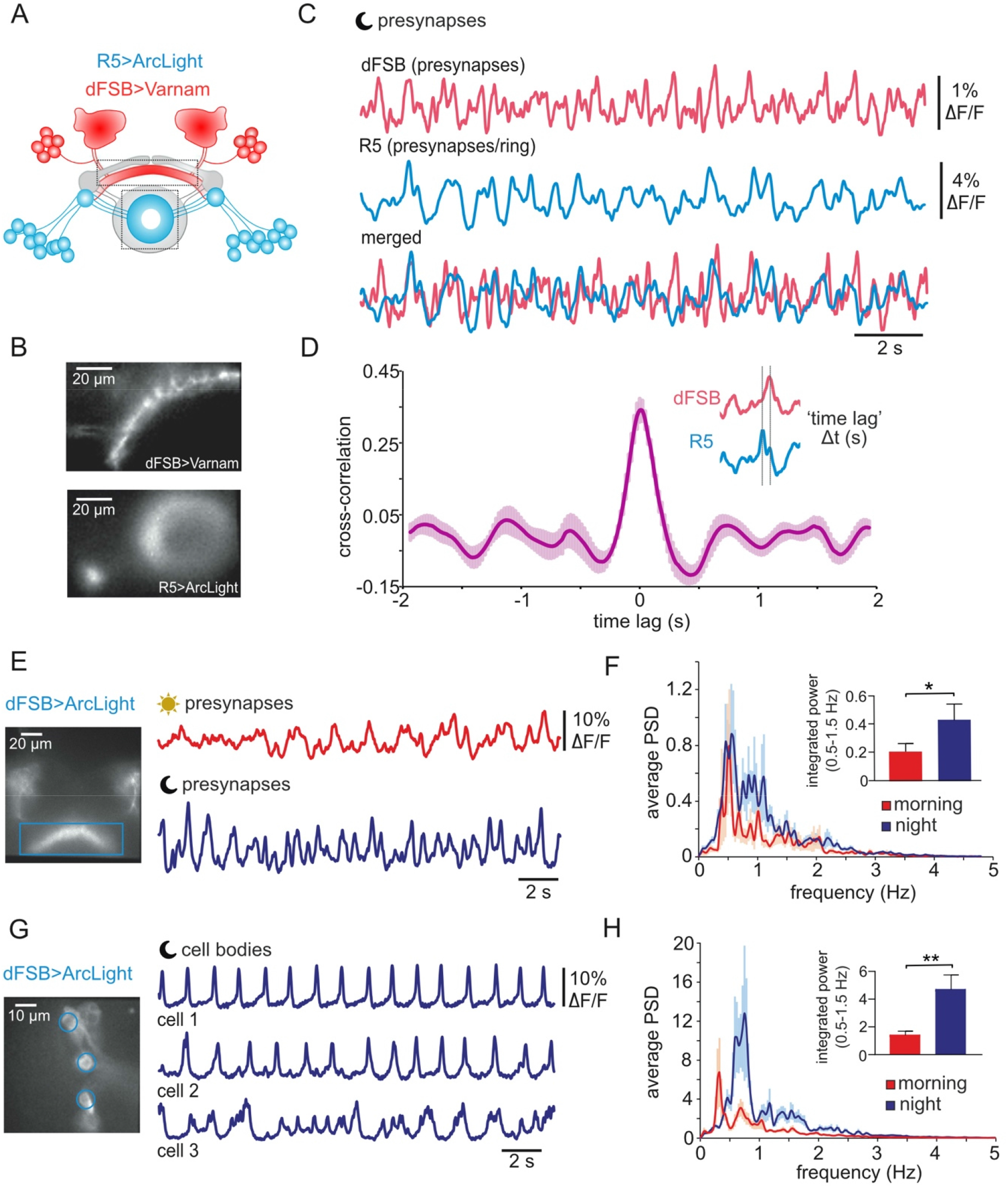
Neural circuits mediating sleep drive display slow-wave coherence. **(A)** Schematic of a sleep regulating circuitry composed of dorsal fan-shaped body (dFSB, red), helicon cells (grey) and R5 neurons (blue). dFSB and R5 neurons are highlighted to indicate expression of genetically encoded voltage indicators (GEVIs). Rectangles indicate recording sites. **(B)** Wide-field image of dFSB neurons expressing Varnam (23E10-LexA) and R5 neurons simultaneously expressing ArcLight (88F06-GAL4). **(C)** Example recording of R5 and dFSB electrical patterns recorded at night (ZT 14-15). **(D)** Cross correlogram of electrical patterns of R5 and dFSB (*n*=10) indicates an overlap of electrical patterns. **(E)** Wide-field image of dFSB neurites and compound recordings of presynaptic electrical activity recorded in the morning (ZT 2-4) and at night (ZT 14-15). **(F)** Average power spectra of compound oscillations in dFSB presynapses. Compound integrated slow-wave power (0.5-1.5 Hz) is increased at night (*n*=9-10, Mann-Whitney test). **(G)** Multi-cellular voltage imaging of individual dFSB neurons recorded in the morning (ZT 2-4) and at night (ZT 14-15). **(H)** Average power spectrum of single-cell voltage recordings showing increased slow-wave power (0.5-1.5 Hz) at night (*n*=23 for ZT 2-4 and *n*=34 for ZT 14-15, Mann-Whitney test).

We conclude that both dFSB and R5 networks exhibit diurnal variations in oscillatory activity and are likely functionally interconnected, as previously reported R5 to dFSB interactions with respect to mediating sleep drive would suggest (Liu et al., 2016).

### Reciprocal functional interactions between neural circuits promoting sleep

To test for functional interactions of network activity patterns, we first asked whether R5 activation could induce dFSB oscillations (**Fig. 2A-C**). We suppressed spontaneous network activity using a previously established high Mg^2+^ extracellular saline (Raccuglia et al., 2019) and optogenetically stimulated R5 neurons using genetic targeting of the red-light shifted channelrhodopsin CsChrimson (Klapoetke et al., 2014) while monitoring intracellular Ca^2+^ in presynaptic terminals of the dFSB (**Fig. 2A**). As presynaptic Ca^2+^ is an adequate proxy for synaptic output, we here focused on compound Ca^2+^ oscillations of the sleep drive-mediating neural circuits. Optogenetic stimulation of R5 reliably induced or amplified dFSB presynaptic SWA between 0-1 Hz (**Fig. 2B, Fig. S2A, B**). This effect was statistically significant only after terminating optogenetic activation (**Fig. 2C**), indicating that activation of R5 neurons can lead to longer lasting physiological changes in the dFSB. This finding is in accordance with the observation that sleep need increases mainly after thermogenetic activation of R5 and moreover requires synaptic output from the dFSB (Liu et al., 2016). Interestingly, the power of the observed SWA in dFSB neurons was more pronounced during the subjective night (**Fig. 2C**), demonstrating the influence of the circadian clock on interactions between sleep drive mediating neural circuits.

**Figure 2:**
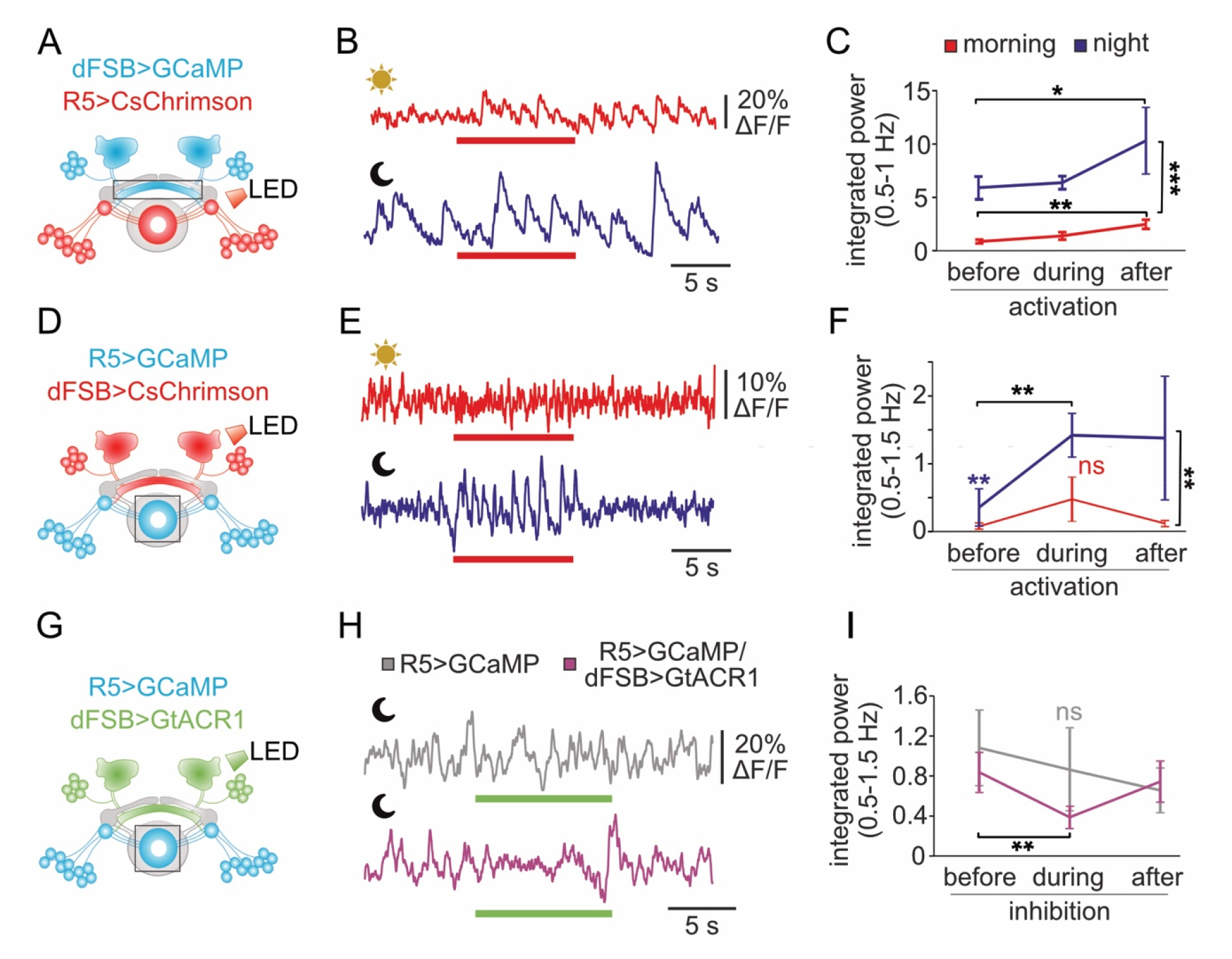
Reciprocal facilitation of SWA between sleep drive mediating circuits is modulated by the circadian clock. **(A, D, G)** Schematic overview of presynaptic recording sites and sites of optogenetic activation and inhibition. **(B)** Example traces of dFSB presynaptic Ca^2+^ activity (23E10-GAL4) during optogenetic activation (red bars) of R5 neurons (58H05-LexA) in the morning (ZT 2-6) and at night (ZT 14-18). **(C)** Slow-wave power (0.5-1 Hz) in the dFSB increases after optogenetic stimulation of R5 neurons (*n*=10, Friedman test with post hoc Dunn’s multiple comparisons test). Pooling the data for during and after stimulation indicates that inducing oscillations is facilitated at night (*n*=20, Mann-Whitney test). **(E)** Example traces of R5 presynaptic Ca^2+^ activity (58H05-GAL4) during optogenetic activation of dFSB neurons (23E10-LexA) in the morning (ZT 2-6) and at night (ZT 14-18). **(F)** At night but not in the morning R5 slow-wave power (0.5-1.5 Hz) increases during optogenetic stimulation of R5 neurons (*n*=10-11, Friedman test with post hoc Dunn’s multiple comparisons test). Pooling the data for during and after stimulation indicates that the induction of oscillations is facilitated at night (*n*=20-22, Mann-Whitney test). **(H)** Example traces of R5 presynaptic Ca^2+^ activity (58H05-LexA) during optogenetic inhibition of dFSB neurons (23E10-GAL4) at nighttime. **(I)** Slow-wave power (0.5-1.5 Hz) is reduced when hyperpolarizing dFSB neurons (*n*=6-7, Friedman test with post hoc Dunn’s multiple comparisons test).

Next, we optogenetically activated the dFSB and recorded presynaptic Ca^2+^ activity in R5 (**Fig. 2D**). Optogenetic activation of the dFSB did not elicit Ca^2+^ oscillations during the day (**Fig. 2E**) but elicited pronounced oscillations between 0.5-1.5 Hz during stimulation at nighttime (**Fig. 2F** and **Fig. S2C, D**). Moreover, in some cases, R5 SWA persisted even after optogenetic activation ceased (**Fig. S2E**), suggesting that increased excitability of the dFSB at night can entrain R5 oscillations and might thus affect sleep need on a larger time scale.

To directly test whether the increased excitability of the dFSB at night drives SWA in R5, we optogenetically hyperpolarized the dFSB using the light-gated chloride channel GtACR (Mohammad et al., 2017) (**Fig. 2G**). To access the potential effect of inhibition, we used a low Mg^2+^ extracellular saline that does not suppress spontaneous activity (Raccuglia et al., 2019). Indeed, activating GtACR in the dFSB reduced oscillatory power of the R5 network (**Fig. 2H, I**), further supporting the notion that sleep need mediating SWA in R5 neurons is facilitated by increased excitability within the sleep-executing neurons of the dFSB. This effect was not observable in controls not expressing GtACR (**Fig. 2H, I**). However, often R5 SWA partially recovered after the start of optogenetic inhibition (**Fig. S2F, G**), suggesting that R5 SWA is stabilized either through other synaptic inputs (such as TuBu) (Lamaze et al., 2018; Raccuglia et al., 2019) or through cell-autonomous promotion of SWA.

Our data so far suggest that functional interactions between network activity in the dFSB and R5 generate sleep drive mediating-SWA dependent on the time of the day. Oscillatory frequencies observed after activating specific networks slightly vary from spontaneously occurring SWA at night, suggesting that compound SWA in the *Drosophila* brain is fine-tuned by an interplay of several networks. After demonstrating that prominent sleep-regulating structures exhibit similar, but not identical mutually interactive oscillations at night, we next addressed to what extent the temporal structure of network activity was relevant for behavior.

### Neural circuits mediating sleep drive acutely control locomotion, postural control and behavioral programs associated with quiescence

Activation of both R5 and dFSB has been shown to have profound impact on fly sleep behavior (Liu et al., 2016; Pimentel et al., 2016; Raccuglia et al., 2019). We now tested how the temporal architecture of electrical patterns that are characteristic of the “sleepy” brain state at night affects behavioral activity during the day.

We devised a paradigm for tracking freely moving flies in a small circular arena while performing temporally fine-tuned optogenetic activation experiments (**Fig. 3A**). To demonstrate that the locomotion of flies in this arena is subject to their internal sleep need, we compared locomotor activity of rested and sleep deprived flies (**Fig. 3B**). As expected, the mean velocity of sleep deprived flies was significantly reduced compared to rested flies (**Fig. 3B**). Moreover, about 56% of sleep deprived flies were immobile for 5 min, which represents the generally accepted criteria for sleep (Shaw et al., 2000), demonstrating that the flies’ locomotor activity in the arena is indeed subject to their internal sleep need.

**Figure 3.**
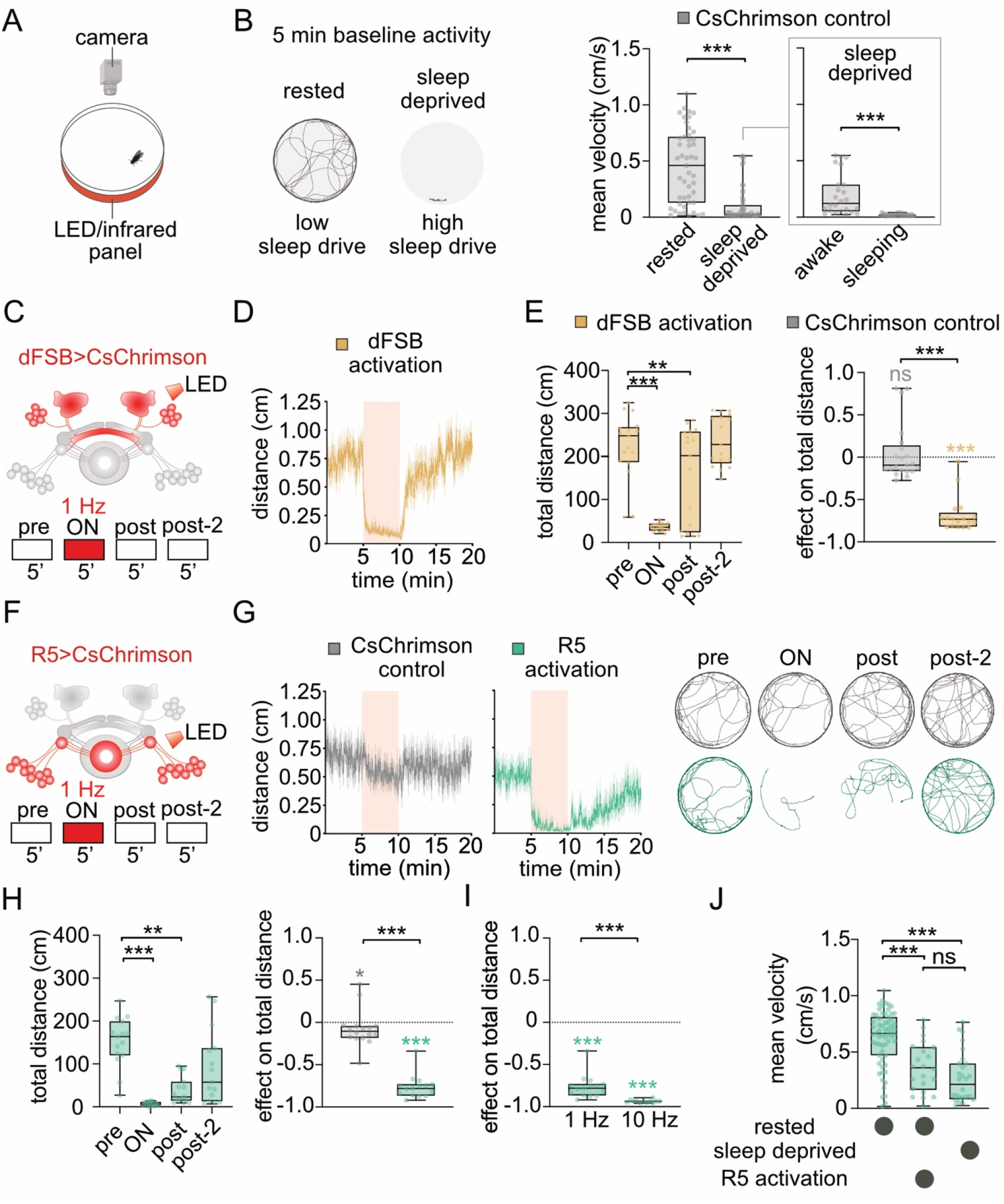
Neural circuits mediating sleep drive acutely control locomotor behavior. **(A)** Locomotor activity is assessed by continuously tracking the position of a single fly in a circular arena. **(B)** Locomotor activity of individual flies correlates to the internal sleep drive. Left: example trajectories (5 min) of a rested and a sleep deprived fly (+>LexAop-CsChrimson). Right: baseline velocity of sleep-deprived flies is reduced compared to that of rested flies (*n*=43 for rested and *n*=55 for sleep deprived, Mann-Whitney test). Sleep deprived flies either meet the sleep criteria (5 min of inactivity, ‘sleeping’, *n*=31) or are active (‘awake’, *n*=24). **(C)** Schematic overview of the optogenetic protocol used to activate dFSB under 23E10-GAL4 control. After 5 min of baseline activity (‘pre’), rested flies were exposed to 5 min red light at 1 Hz (‘ON’) and allowed to recover for 10 min (‘post’ and ‘post-2’). **(D)** Average locomotion trace of 23E10-GAL4>UAS-CsChrimson flies (+ retinal, *n*=17). Curves: mean ± SEM. **(E)** Left: 1 Hz dFSB activation decreases locomotion of rested flies during and after activation (*n*=17, Friedman test with Dunn’s multiple comparisons). Right: optogenetic stimulation of the dFSB (+ retinal) reduced total distance travelled compared to control flies (+ retinal), *n*=17-19, Mann-Whitney test (comparisons between genotypes depicted in black) and Wilcoxon signed-rank test (comparisons to baseline locomotion depicted in color). **(F)** The same optogenetic protocol described in **C** is used to activate R5 neurons under 58H05-GAL4 control. **(G)** Left: average locomotion traces of CsChrimson control (+ retinal, *n*=16) and 58H05-GAL4>UAS-CsChrimson flies (+ retinal, *n*=15). Curves: mean ± SEM. Right: example trajectories. **(H)** Left: 1 Hz stimulation reduced distance travelled during and after activation of R5 (*n*=15, Friedman test with Dunn’s multiple comparisons). Right: optogenetic activation of R5 decreased distance travelled compared to control flies, *n*=15-16 each group, Mann-Whitney test (comparisons between genotypes depicted in black) and Wilcoxon signed-rank test (comparisons to baseline locomotion depicted in color). **(I)** The effect of R5 stimulation on distance travelled differed between different activation frequencies (*n*=15 for 1 Hz and *n*=12 for 10 Hz, Mann-Whitney test). **(J)** Mean walking velocity was reduced when R5 was activated at 1 Hz, resembling the locomotion levels of awake but sleep-deprived flies (+ retinal) (*n*=63 for rested, *n*=22 for rested+R5 activation, *n*=28 for sleep deprived, Kruskal-Wallis test with Dunn’s multiple comparisons). Experiments shown in this figure were performed during the subjective day of the flies (ZT 1-10).

We first asked whether 1 Hz optogenetic activation of the dFSB could acutely regulate locomotor activity (**Fig. 3C**). Importantly, optogenetic activation of dFSB neurons at 1 Hz had previously been shown to reliably promote the naturally occurring oscillatory pattern in single dFSB neurons (Troup et al., 2018). To be more sensitive to the induction of potential sleep states, we performed these experiments during the subjective day (ZT 1-10). 1 Hz stimulation of the dFSB decreased the walking velocity as well as the time spent walking (**Fig. 3D-E** and **S3A**), which is in accordance with the strongly sleep-promoting function of the dFSB. To exclude the possibility that the observed behavior was a visually triggered response to pulsed light, we evaluated the effect of the optogenetic protocol on the locomotion of flies carrying only the UAS-CsChrimson transgene. We found that pulsed light slightly reduced locomotor activity (**Fig. 3E, S3A**, compare to **3G**). However, inhibition of locomotor activity through dFSB activation was considerably stronger (**Fig. 3E** and **S3A**). To test whether inhibition of locomotion was frequency-specific, we next stimulated dFSB neurons at 10 Hz. We observed locomotor phenotypes comparable to 1 Hz stimulation, indicating that the activation frequency and correlating increase in stimulation intensity was not crucial to increase sleep need (**Fig. S3B**).

We next tested whether acute rhythmic optogenetic activation of R5 would similarly affect locomotion (**Fig. 3F**). Indeed, activating R5 at 1 Hz significantly reduced locomotion similar to what we observed when stimulating the dFSB (**Fig. 3G, H, S3C**). During optogenetic stimulation of R5, flies displayed short walking bouts as well as frequent grooming (**Fig. S3F**). Unlike what was observed when stimulating the dFSB, locomotion recovered much slower after stimulation offset, not reverting to normal for the following 5 min (**Fig. 3G, H**). This is in line with R5 activity entraining the involved neural circuits to prepare flies for sleep and promote more consolidated sleep phases (Raccuglia et al., 2019). To determine whether flies still showed appropriate locomotor behavior during optogenetic stimulation of R5, we investigated whether flies were capable of reaching higher velocities while walking (“walking velocity”) by only considering movements > 0.25 cm/s (see **Methods**, **Fig. S3C**). We found that this walking velocity was only slightly affected (**Fig. S3C**), indicating that reduced locomotion was mainly due to a reduction in time spent walking. Moreover, we also examined the flies’ behavioral responsiveness to an air-puff during 1 Hz R5 stimulation and found that flies reacted to this strong stimulus by increasing their walking velocity (**Fig. S3D**). These findings suggest that R5 activation at 1 Hz reduces locomotor activity while leaving the ability to walk and respond to strong stimuli intact. We therefore conclude that 1 Hz R5 activation induces a quiescent state characterized by reduced locomotor activity and frequent grooming, which in fact has been associated with periods of rest that occur prior to sleep (Qiao et al., 2018).

We next explored whether the behavioral outcome following R5 activation was frequency-specific. Unlike the effects observed when stimulating at 1Hz, 10 Hz protocols rendered flies immobile and ∼90% of flies lost postural control during optogenetic activation, sometimes lasting for 5 min post-stimulation (**Fig. 3I, S3E-G**). Stimulating R5 neurons at 0.1 Hz did not lead to a measurable impact on the flies’ behavior (**Fig. S3E**). Therefore, contrary to the dFSB, the activation frequency and intensity mattered for the R5 network (**Fig. 3I and S3E**), potentially placing the R5 network as instructive for an ongoing brain state (such as sleep). While the slow oscillations observed in R5 allows the flies to maintain natural body posture during quiescence, it is possible that over-stimulating R5 neurons at 10 Hz overrules downstream effectors involved in standing upright. In line with this, 1 Hz optogenetic activation of R5 in rested flies reduced locomotor activity to the level of awake sleep deprived flies (**Fig. 3J**), indeed suggesting that 1 Hz R5 activation is specific to increasing sleep need in awake animals. Taken together, these data indicate that acute activation of neural circuits mediating sleep drive at a frequency band of sleep-relevant SWA induces a quiescent state characterized by reduced locomotion.

### Sleep-promoting R5 neurons and locomotion-gating helicon cells synchronize electrical patterns at night

Our data demonstrate reciprocal functional interactions between the dFSB and R5 networks and also indicate that these structures can act as functional analogs at the behavioral level. Using the recently published hemiconnectome of *Drosophila* (Scheffer et al., 2020), we next investigated putative neural connections between the dFSB and R5 and found no direct synaptic connections (**Fig. 4A, B**). Instead, our analysis revealed that R5 neurons are highly interconnected with the previously identified helicon network (Donlea et al., 2018). These four neurons make reciprocal all-to-all synaptic connections with R5 (**Fig. S4A**) that represent about 24% of R5 input and output connections (**Fig. 4A**). Similarly, R5 neurons are one of the main pre- and postsynaptic partners of helicon cells (**Fig. S4B**). Consistent with previous observations showing that dFSB neurons inhibit visually evoked responses in helicon cells (Donlea et al., 2018), we found that helicon cells are reciprocally connected to the dFSB (**Fig. S4C**) (Hulse et al., 2021). Our analysis further indicates that, as previously suggested (Donlea et al., 2018), recurrent connections from helicon cells could constitute a main route to relay neural information between the dFSB and R5 networks and vice versa (**Fig. 4B and S4D**).

**Figure 4.**
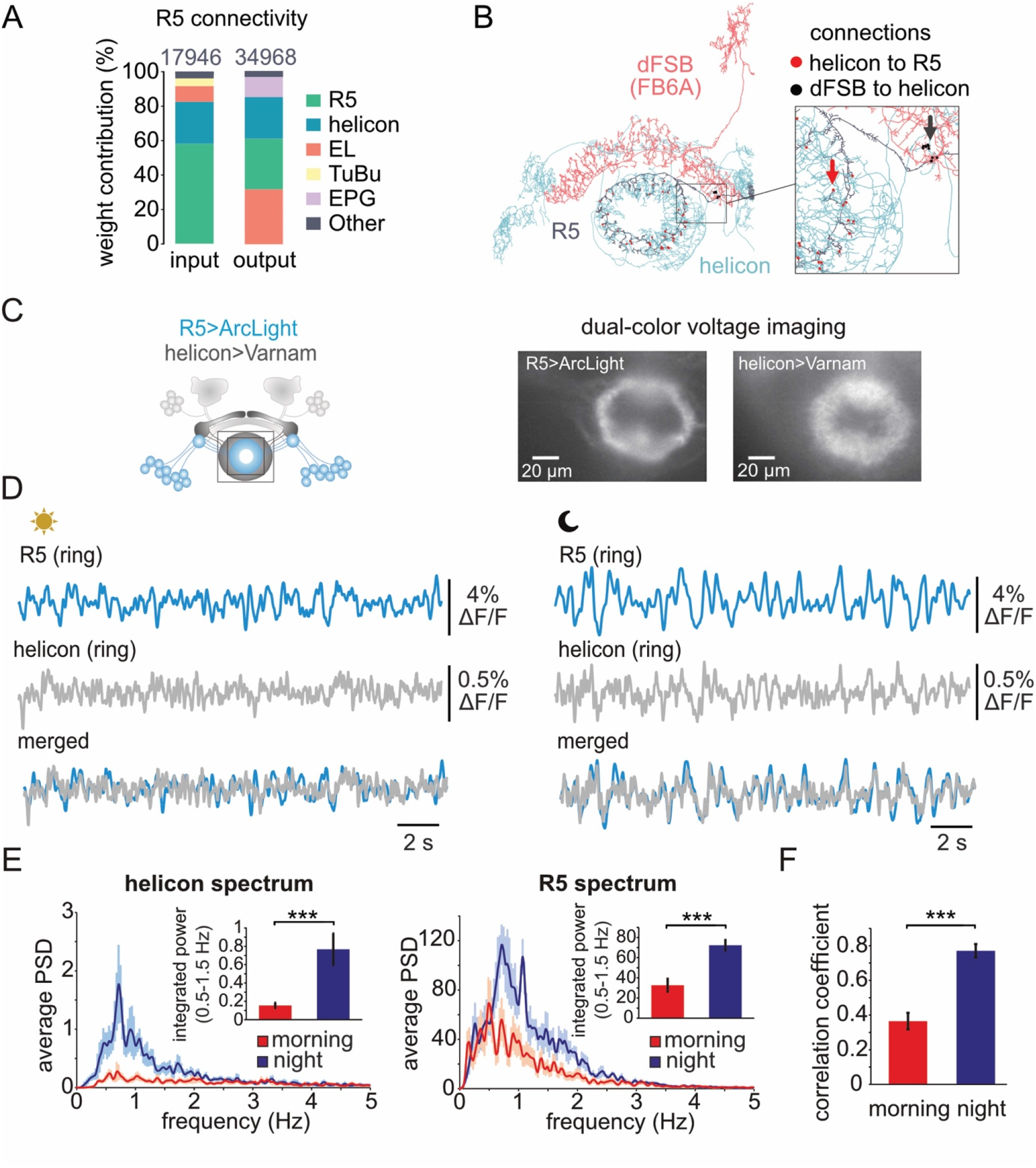
Sleep drive mediating and arousal gating networks synchronize electrical slow-wave patterns at night. **(A)** Synaptic input and output connections from R5 neurons show high connectivity with helicon cells. **(B)** Example circuit between a R5 neuron (body ID: 1200049187), helicon cell (body ID: 918334668) and dFSB neuron (neuron type FB6A) (body ID: 422191200). Red dots/arrow indicate areas of synaptic connections from the helicon cell to the R5 neuron and black dots/arrow areas of synaptic connections from the dFSB neuron to the helicon cell. **(C)** Schematic and wide-field image of presynaptic ring structures expressing the green voltage indicator ArcLight in R5 neurons (88F06-GAL4) and the red voltage indicator Varnam in helicon cells (24B11-LexA). Square indicates recording site. **(D)** Example recordings of R5 and helicon presynaptic electrical patterns recorded in the morning (ZT 2-4) and at night (ZT 14-16). **(E)** Average power spectra of compound oscillations in R5 neurons and helicon cells demonstrate that slow-wave power (0.5-1.5 Hz) is increased at night in both networks (*n*=9-11 for helicon cells and R5 neurons, Mann-Whitney test). **(F)** Correlation between electrical patterns of helicon and R5 networks is significantly increased at night compared to the subjective morning (*n*=9-11, Mann-Whitney test).

Since helicon cells and R5 neurons seem to play a role in both visually guided navigation and mediating sleep need (Flores-Valle et al., 2021; Seelig and Jayaraman, 2013), it is likely that temporal interactions between the activity patterns of these circuits regulate locomotor activity during wake and sleep. To first probe for electrical network activity in helicon cells at night, we expressed the voltage indicator ArcLight (**Fig. S4E**) and found spontaneous compound activity between 0.5-1.5 Hz (**Fig. S4F, G**), indicating similarities to R5 oscillatory activity.

To investigate a direct relationship between R5 and helicon electrical patterns, we simultaneously recorded the electrical compound activity at their overlapping presynaptic sites by performing dual color voltage imaging experiments expressing the green voltage indicator ArcLight in R5 and the red voltage indicator Varnam in helicon (**Fig. 4C**). We confirmed that in both R5 and helicon oscillatory power (0.5-1.5 Hz) is increased at night compared to the subjective morning (**Fig. 4D, E**). Most interestingly, correlation between the electrical patterns of these neural populations was significantly increased at night (**Fig. 4F**). To control for different kinetics of the voltage indicators, we swapped the expression (expressing ArcLight in helicon cells and Varnam in R5 neurons) and observed similarly high correlation at nighttime (**Fig. S4H-J**). Therefore, our data suggest that helicon compound activity might be entrained by R5 slow-wave activity.

### Modelling predicts that R5 entrains helicon oscillation via excitatory connections

Based on our physiological findings, we next turned to theoretical modeling to further investigate how network interactions could steer behavior. Our data suggests that R5 could function as the master oscillator, entraining synchronous oscillatory activity in helicon. We used the Izhikevich model to simulate synchronization of single R5 neurons (please see **Supplemental text** and **Fig. S5-1** legend for detailed information). R5 neurons are highly interconnected (**Fig. S5-1A**), which could allow them to synchronize through mutual interaction. R5 neurons, however, have been identified to be GABAergic (Hanesch et al., 1989), a finding we confirmed using immunofluorescence (**Fig. S5-1C**). Not surprisingly, as inhibitory pulse-coupled oscillators are known to usually stay out of sync (Terman et al., 1998), we found that in order to simulate synchronization patterns similar to those experimentally observed, we required mixed excitatory and inhibitory output of R5 neurons (**Fig. S5-1E**). In the model, we varied the relative strength of the mixed inhibitory and excitatory synaptic coupling among R5 neurons and observed that synchronization correlated with the relative strength of excitation compared to inhibition within the network (**Fig. S5-1E-H**). The main excitatory neurotransmitter in insect brains is acetylcholine (Fayyazuddin et al., 2006). To experimentally verify whether R5 also had excitatory outputs, we used an endogenously HA-tagged vesicular acetylcholine transporter (VAChT), a bona fide marker for cholinergic vesicles, that can be induced to label cholinergic output sites in specific neural populations (Pankova and Borst, 2017). Strikingly we identified several cholinergic R5 sites (**Fig. S5-1B**), supporting a R5 network with mixed inhibitory and excitatory output as source for the observed oscillations. We next added helicon cells to our model to investigate coherent activity of helicon and R5 networks throughout the day. Helicon, as a purely excitatory network recurrently connected to R5, was modeled to switch from a more polarized and active state in the morning (‘upstate’) to a more hyperpolarized, inactive state (‘downstate’) at nighttime (**Fig. 5A**). This was warranted, as dFSB stimulation had previously been shown to hyperpolarize helicon (Donlea et al., 2018). Moreover, providing further experimental data for our model, we found through baseline fluorescence quantification of the voltage indicator ArcLight, that single helicon cells were more hyperpolarized in the evening compared to the morning hours (**Fig. S5-1D**). Therefore, our simulation is in line with the dFSB paving the path for R5-mediated association of helicon activity at night (compare **Fig. 2**) by rendering helicon less responsive to inhibitory (while still responsive to excitatory) input from R5 when helicon’s resting membrane potential is closer to the chloride reversal potential (Barker and Harrison, 1988). Indeed, our simulation resulted in 1 Hz oscillations at night for both helicon and R5, while helicon was not oscillating in the daytime settings (**Fig. 5B**). Moreover, reminiscent of our experimental findings (**Fig. 4F**), both networks were highly synchronized at night and R5 activity slightly preceded helicon activity, suggesting that R5 drives compound oscillations in helicon cells (**Fig. 5C**). Given the close match between our simulation and experimental findings, we used our model to predict the outcome of experiments performing artificial stimulations. To do so, we simulated network activity when stimulating either network or both together (**Fig. 5D**). Strikingly, we found that the underlying network activity architecture showed dominant R5 activity over helicon (**Fig. 5E**), suggesting that activation of R5 and helicon at the same time should have a similar behavioral outcome as activating R5 on its own (please see supplemental information for more details). Indeed, simultaneous activation closely resembles the observed nighttime configuration (**Fig. 5B**). We turned back to behavioral experiments to test this prediction.

**Figure 5.**
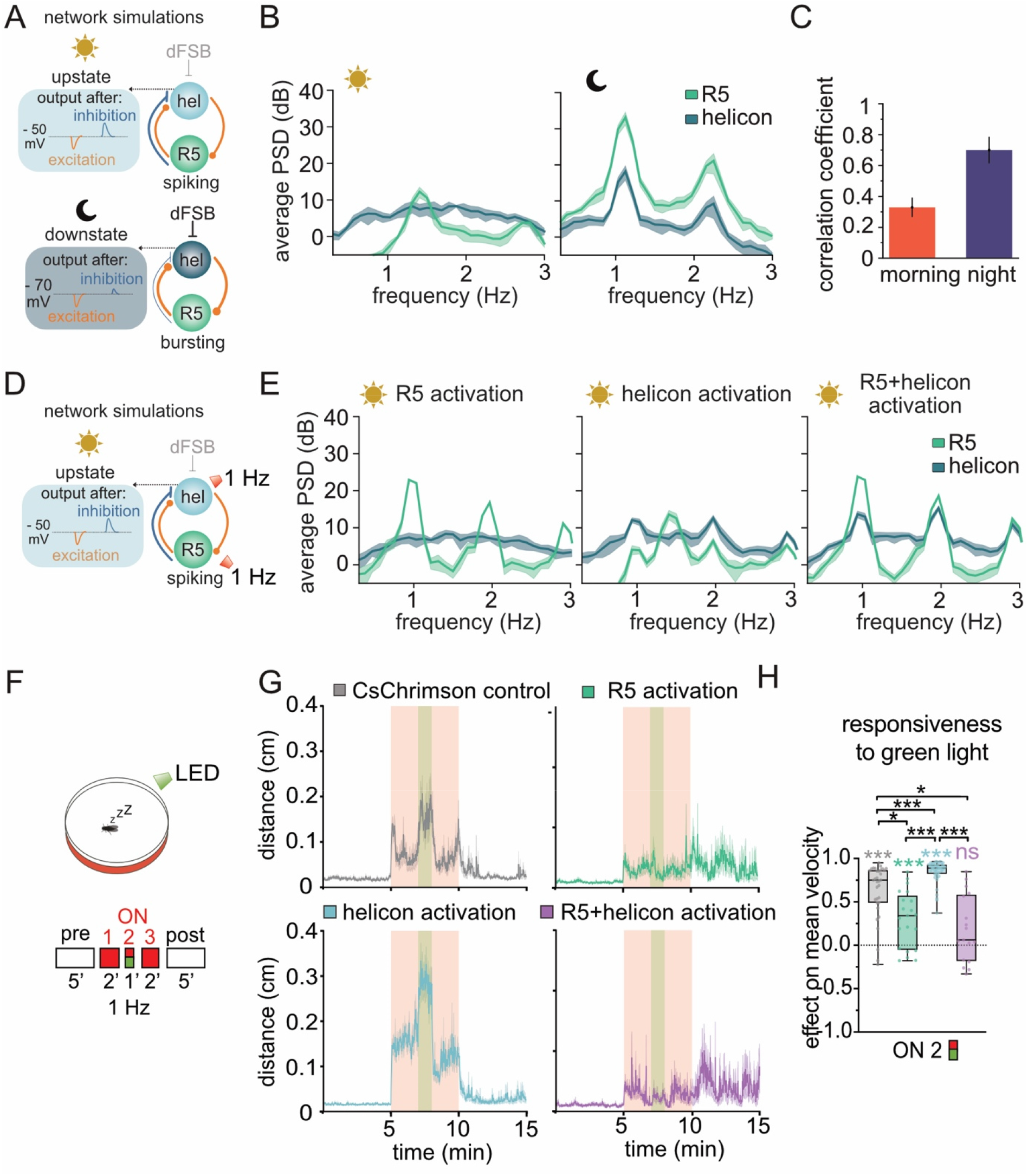
R5 entrains coherent oscillations in helicon to regulate behavioral responsiveness. **(A)** Schematic overview of simulated R5 and helicon interactions in the morning (top) and night (bottom). In our model, the helicon network is purely excitatory while R5 is mixed inhibitory and excitatory (see Fig. S5-1). In the morning configuration, R5 neurons spike and helicon cells are in a depolarized state (‘upstate’). In the night configuration, R5 neurons burst, and due to inhibition mediated by the dFSB helicon cells are hyperpolarized (‘downstate’). Please see Fig. S5-1 and Supplemental Experimental Procedures for more detailed information. **(B)** Simulation of R5 and helicon synchronization during the day and during the night. R5 was coupled to helicon to assess levels of synchronization within and between networks. Power spectral density (PSD) of the R5 and helicon networks in the morning and at night. In the morning, R5 did not entrain helicon activity. At night, R5 neurons became highly synchronized and entrained helicon activity to a 1 Hz rhythm, which further promoted R5 synchronization (n=10 simulations each). **(C)** Simulated correlation coefficients of compound R5 and helicon activity in the morning and at night. Similar to the experimental data (Fig. 4F), correlation of the compound activity is increased at night (n=10 simulations each). **(D)** To predict the outcome of network interactions, R5 and helicon were individually and simultaneously stimulated with 1 Hz in the morning configuration. **(E)** Average PSD of R5 and helicon networks during 1 Hz stimulation in the morning configuration. 1 Hz R5 stimulation led to R5 synchronization and no helicon entrainment. 1 Hz stimulation of the helicon network led to a slight entrainment of helicon to a 1 Hz rhythm and no R5 synchronization. Simultaneous stimulation of R5 and helicon at 1 Hz led to strong R5 synchronization and helicon entrainment (n=10 simulations). **(F)** Schematic overview of optogenetic activation protocol testing behavioral responsiveness in sleeping flies. After 5 min of immobility, sleeping flies were presented with 5 min of red light at 1 Hz and a continuous green light during the stimulation. **(G)** Average locomotion traces of CsChrimson control (+ retinal, *n=33*), 58H05-LexA>LexAop-CsChrimson (R5; + retinal, *n*=22), 24B11-LexA>LexAop-CsChrimson (helicon, + retinal, *n=47*) and 58H05-LexA; 24B11-LexA>LexAop-CsChrimson flies (R5 + helicon; + retinal, *n=17)*. Curves: mean ± SEM. **(H)** Behavioral responsiveness to green light during optogenetic activation (colored asterisks indicate significance compared to baseline locomotion). Behavioral responsiveness during R5 activation was reduced compared to helicon stimulation and control. Simultaneous activation of R5 of helicon led to the strongest impairment in behavioral responsiveness, *n*=17-47 each group, Kruskal-Wallis test with Dunn’s multiple comparisons (black asterisks). Behavioral data was taken during the subjective day of the flies (ZT 1-7) after a 12-19 h of sleep deprivation.

### Network coherence regulates behavioral responsiveness

In mammals, synchronization of SWA between and within neural circuits occurs during deep sleep and local sleep in awake individuals and is hypothesized to establish neural filters that suppresses behavioral responsiveness to external stimuli (Quercia et al., 2018). To investigate whether synchronization between R5 and helicon play a role in regulating behavioral responsiveness during sleep, we used optogenetics to separately and simultaneously activate these networks at 1 Hz in sleeping flies (**Fig. 5F-H**). After sleep deprivation, sleeping flies were exposed to green light to arouse them and elicit visually evoked locomotion while simultaneously applying red light pulses at 1 Hz for network-specific optogenetic stimulation (**Fig. 5F**). As shown for rested control flies (compare **Fig. 3G**), all genotypes showed slight arousal to red light (**Fig. 5G, and S5-2A, B**). However, as expected, helicon cell stimulation led to the strongest degree of arousal (**Fig. S5-2A**) with most flies being awakened (**Fig. S5-2B**), while arousal during R5 stimulation was consistent with the control (**Fig. S5I**). During stimulation with green light, control flies and helicon activation showed the highest degree of visually induced locomotion (**Fig. 5G-H**). In contrast, visually triggered locomotion was not observed during stimulation of R5 neurons. While flies were capable of reacting to a strong air-puff (**Fig. S3D**), optogenetically induced R5 oscillations seem to establish a neural filter that blocks visually induced locomotion. Interestingly, compared to R5 activation alone, synchronous activation of R5 and helicon led to a stronger reduction of locomotor activity (**Fig. 5H**) and only a small fraction of flies were awakened by green light (**Fig S5-2B**), indicating that R5 and helicon synchronization might establish a more efficient neural filter. Together, these findings indicate that sleep drive mediated by R5 oscillations overrides gating of locomotion by helicon cells and further demonstrates that synchronization between these networks locks the locomotion-initiating networks into a state that, dependent on sleep need, reduces sensory processing and behavioral responsiveness. Of note, in the model, R5 and helicon activation led to a slightly larger 1 Hz oscillation of the R5 network than the activation of R5 alone (**Fig. 5E**). Similarly, a 1 Hz activation of only helicon cells had a relatively small impact on inducing oscillations in R5 (**Fig. 5E**), which matches the behavioral observation that helicon cell stimulation induced arousal and not quiescence (**Fig. 5G** and **Fig. S5-2A**).

### Helicon and R5 antagonistically regulate locomotor activity and behavioral responsiveness via modulation of head direction neurons

We next aimed at investigating the mechanistic basis of how R5 and helicon synchronization could regulate behavioral responsiveness. We therefore turned back to the connectome and found that helicon cells as well as R5 neurons are connected to EPG neurons (**Fig. 6A**), which are head-direction neurons representing the fly’s heading direction and initiating body turns towards or away from sensory stimuli (Green et al., 2017; Hulse et al., 2021; Kim et al., 2017; Turner-Evans et al., 2017; Vafidis et al., 2022). In fact, we found that for about 70% of R5-to-EPG synaptic connections the nearest non-R5 neighbor were helicon-to-EPG synapses (**Fig. 6A, B**). Moreover, within the inner parts of the ring, where there are no R5 synapses, helicon-EPG synapses can be found in close proximity to synaptic input of sleep-regulating R3 neurons (Aleman et al., 2021; Hulse et al., 2021) (**Fig. 6B**), indicating that these synaptic motifs and their spatial relations could be major sites of integrating navigation and sleep regulation (Flores-Valle et al., 2021).

**Figure 6:**
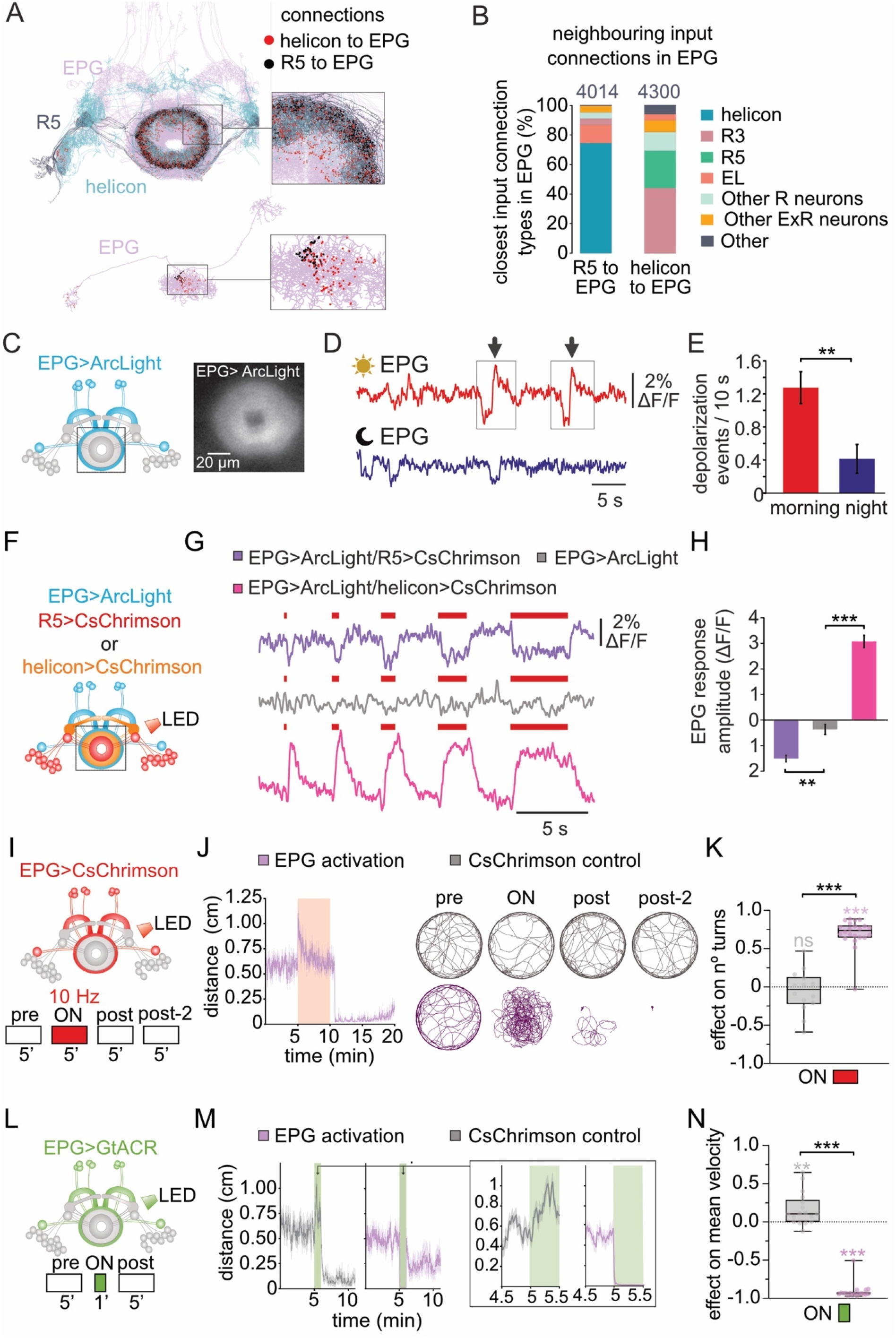
R5 and helicon regulate behavioral responsiveness via antagonistic inputs to the navigational system. **(A)** Top: the helicon-R5-EPG circuitries are strongly interconnected at the level of the ring. Morphological renderings of R5 neurons, helicon cells and EPG neurons. Red dots depict synaptic connections of helicon cells to EPG neurons and black dots show synaptic connections of R5 to EPG. Bottom: Location of R5 and helicon synapses projecting to a single EPG neuron (bodyId: 387364605). **(B)** At the level of EPG neuron dendrites, synaptic input connections from R5 neurons (R5-to-EPG) are found in proximity to input connections from helicon cells (helicon-to-EPG). Similarly, helicon-to-EPG synaptic connections are found in proximity to R5- to-EPG connections and other ring neurons-to-EPG connections. Bar plots indicate the fraction of closest synaptic connections for R5-to-EPG and helicon-to-EPG per neuron type. The numbers of R5-to-EPG and helicon-to-EPG synaptic connections analyzed are indicated. **(C)** Schematic overview and wide-field image of EPG neurons expressing the voltage indicator ArcLight (60D05-GAL4). **(D)** Example recordings of EPG electrical patterns recorded in the morning (ZT 2-4) and at night (ZT 14-16) indicating reduced activity at night. Squares indicate spontaneous non-oscillatory events during the day. **(E)** Incidence of depolarization events is markedly reduced at night (*n*=12, Mann-Whitney test). **(F)** Schematic overview of EPG recording site (60D05-GAL4) during optogenetic activation of R5 (58H05-LexA) and helicon (24B11-LexA). **(G)** Example recordings showing that activation of R5 neurons leads to net hyperpolarization of the EPG network and that helicon cells depolarize it. To suppress spontaneous activity during helicon activation we used a high Mg^2+^ solution. **(H)** EPG response amplitudes during optogenetic activation reveals that R5 mediates hyperpolarization while helicon mediates depolarizations, n=25-30 stimulations in 5-6 independent brains, Kruskal-Wallis with posthoc Dunn’s test. **(I)** Schematic overview of the optogenetic protocol (as described in Fig. 3C) used to activate EPG neurons under control of 60D05-GAL4. **(J)** Left: average locomotion traces of 60D05-GAL4>UAS-CsChrimson and control flies (+ retinal, *n*=22). Right: example trajectories. **(K)** Optogenetic activation of EPG at 10 Hz increased number of turns, *n*=14-22 each group, Mann-Whitney test (comparisons between genotypes, black asterisks) and Wilcoxon signed-rank test (comparisons to baseline locomotion, colored asterisks). **(L)** Schematic overview of the optogenetic protocol used to hyperpolarize EPG neurons expressing UAS-GtACR1 under the control of 60D05-GAL4. After 5 min of baseline activity, flies are exposed to 1-min continuous green light from below and allowed to recover for 5 min. **(M)** Average locomotion traces of GtACR1 control (+ retinal, *n*=17) and 60D05-GAL4>UAS-GtACR1 flies (+ retinal, *n*=18). Curves: mean ± SEM. **(N)** Constant inactivation of EPG neurons drastically reduced mean velocity in 60D05-GAL4>UAS-GtACR1 flies compared to control flies, *n*=17-18 for each group, Mann-Whitney test (comparisons between genotypes, black asterisks) and Wilcoxon signed-rank test (comparisons to baseline locomotion, colored asterisks).

To test whether activity in the head-direction system is, similar to R5 and helicon, subject to diurnal variation we used ArcLight to record the EPG compound electrical activity (**Fig. 6C-E**). In the morning, EPG neurons displayed non-oscillatory events comprised of de- and hyperpolarizations (**Fig. 6D**). Depolarizations occurred at an average rate of 1.2 every 10 s in the morning and significantly dropped at night (**Fig. 6E**).

By expressing CsChrimson in R5 or helicon we next tested whether these networks could be responsible for regulating activity in the EPG network (**Fig. 6F**). In fact, we found that R5 hyperpolarizes EPG neurons, while activation of helicon cells led to strong depolarizations (**Fig. 6G, H**), indicating that R5 and helicon cells modulate EPG activity via antagonistic neurotransmitter inputs.

Our data therefore predict that de- and hyperpolarizing EPG neurons would have opposing effects on locomotor activity and behavioral responsiveness. We tested this by directly de- and hyperpolarizing EPG in rested flies using CsChrimson and GtACR respectively (**Fig. 6I-N**). While depolarizing EPG neurons at 1 Hz had no effect on locomotion (**Fig. S6A**), activating EPG neurons at higher frequencies (10 Hz) increased locomotor activity particularly at the beginning of the stimulation (**Fig. 6J**). Moreover, flies displayed increased turning behavior throughout the stimulation (**Fig. 6K**). Interestingly, immediately after ceasing EPG activation flies transitioned to a quiescent state with starkly reduced locomotion (**Fig. 6J**), which suggests homeostatic regulation of locomotor activity through EPG neurons.

We next tested whether GtACR-mediated hyperpolarization of EPG can actually reduce locomotor activity (**Fig. 6L**). In control flies, constant green light leads to a sudden increase in locomotor activity (**Fig. 6M, N**), demonstrating the behavioral responsiveness to visual stimuli. When we simultaneously inhibited EPG, flies did not respond to green light but rather completely stopped moving and about 45% of flies lost postural control (**Fig. 6M, N and S6D, E**). This behavioral outcome is reminiscent of R5 activation (**Fig. 3 and 5**), indicating that inhibition of EPG activity mediated through R5 can be a crucial component for fine-tuning navigation during the day as well as modulating behavioral responsiveness and sleep depth during the night. Together, our data is consistent with a mechanism, where R5 overrules input of helicon to EPG at night by temporally associating helicon activity and thus creating a neural filter to promote quiescence.

## Discussion

The occurrence of brain-wide SWA during deep sleep (Adamantidis et al., 2019; Buzsaki and Draguhn, 2004) as well as network-specific SWA during local sleep in awake individuals (Hung et al., 2013; Quercia et al., 2018) suggests that coherent SWA could represent a neural filtering mechanism that regulates sensory processing and behavioral responsiveness. How coherent network activity suppresses behavioral responsiveness to promote a quiescent egocentric state remains unclear to this day. Here, we discover a mechanism in *Drosophila*, where neural networks promoting sleep need generate coherent oscillations across neural networks in order to reduce the functional connectivity between locomotion gating networks and the navigational system.

In this study, we demonstrate, to our knowledge for the first time, that SWA can provide the temporal architecture that creates a “breakable” neural filter by providing an inhibitory tone that suppresses gating of locomotion and at the same time maintains postural control to be able to quickly respond to strong or salient stimuli (**Fig. 7**). Moreover, we reveal that network interactions can form the mechanistic basis of filtering functions by temporally associating antagonistic neurotransmitter inputs to impede behavioral responses to external stimuli (**Fig. 7**).

**Figure 7:**
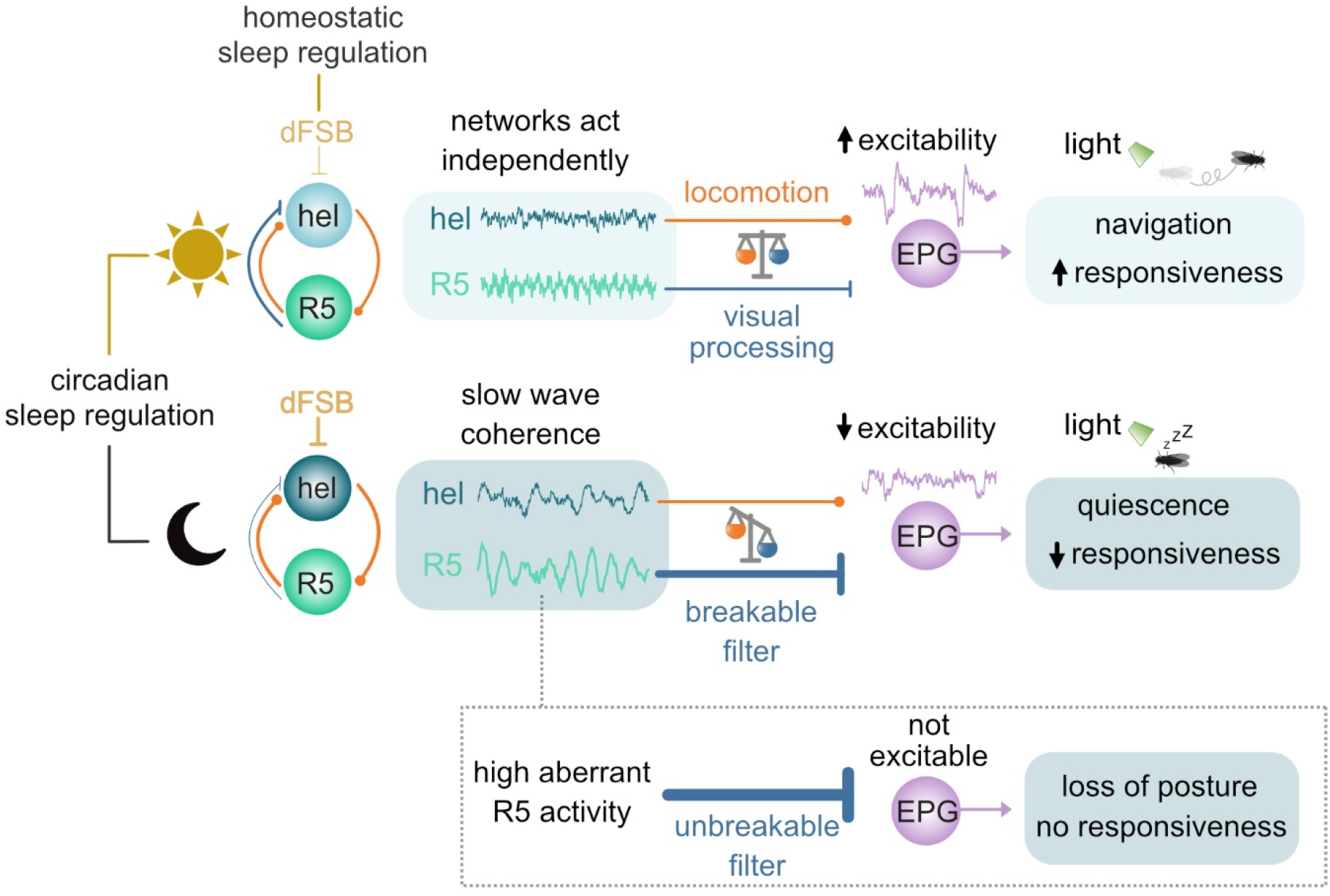
Diurnal modulations of network interactions generate coherent slow-wave activity that act as neural filters to promote quiescence Daytime: Due to reduced activity of the fly’s sleep homeostat (dFSB), R5 does not entrain slow-wave activity in helicon (hel). High frequency output from helicon gates locomotion via excitation (orange line) of head-direction neurons (EPG), which promotes navigation and increases responsiveness to visual stimuli. **Nighttime:** Increased activity of the dFSB and circadian regulation hyperpolarize helicon and drive entrainment of slow-wave coherence through R5. Network coherence and the temporal structure of inhibitory input provided by R5 reduce excitability of EPG neurons, creating a breakable neural filter, which promotes quiescence and reduces behavioral responsiveness. Aberrant high inhibition of EPG by R5 leads to loss of posture and induces a coma-like reversible state of complete unresponsiveness.

### Generating SWA is inherent to neural networks

We discovered that the diurnal cycle generates sleep need that elicits slow oscillatory activity within a previously described recurrent circuitry (Donlea et al., 2018). At night, the master sleep homeostat, the dFSB, increases excitability (Kempf et al., 2019) to generate compound oscillations (**Fig. 1**) which in turn promote partially coherent oscillatory activity in the sleep need mediating R5 network (**Fig. 1, 2**).

Interestingly, we found that dependent on the circadian clock the dFSB and the R5 network reciprocally facilitate presynaptic Ca^2+^ oscillations (**Fig. 2**) and propose the helicon cells (**Fig. 4**) as mediator between the networks. The diurnal variation in functional connectivity is most likely based on the R5 network’s increase in presynaptic output weights when sleep need increases (Liu et al., 2016). The changes in synaptic weight might also promote the nighttime specific synchronization between helicon and R5 networks (**Fig. 4**), which we found to be an essential part of the filtering mechanism that promotes a quiescent egocentric mode (**Fig. 5**).

Similar to neural oscillations in mammalian circuits, we observed slight changes in the power spectrum of spontaneous SWA compared to the SWA induced by activating specific circuits. This is likely explained by the fact that the precise power and frequency of SWA depends on a complex interplay between different neural circuits (Neske, 2015). However, *ex vivo* recordings have shown that even relatively small isolated cortical and thalamic circuits have the ability to generate SWA (Neske, 2015), indicating that neural networks can oscillate autonomously. Moreover, the homeostatic regulation of SWA can occur locally in the brain, indicating that sleep need is experienced by specific networks (Huber et al., 2004). Our findings in *Drosophila* further substantiate the notion that SWA is inherent to neural circuits and that network coherence could represent an evolutionarily optimized strategy to create neural filter mechanisms (Raccuglia et al., 2019; Suárez-Grimalt and Raccuglia, 2021).

### Coherent network oscillations create a filter mechanism to promote quiescence

Our data indicate that the dFSB acts as a switch that promotes oscillations in sleep need mediating R5 (**Fig. 1, 2**) which in turn entrain oscillatory activity in arousal mediating helicon (**Fig. 5**) thus promoting a quiescent state (**Fig. 3**) with reduced behavioral responsiveness (**Fig. 5**).

The observed network interactions have intriguing functional and physiological analogies to thalamocortical interactions in mammalian brains. The thalamus orchestrates cortical synchronization during sleep and functions as a relay station for sensory signals and motor commands, essentially regulating behavioral responsiveness to external stimuli (Andrillon and Kouider, 2020). However, how coherent oscillations can function as a gate remains unclear. Here, our work provides mechanistic insight (**Fig. 7**): during the day, helicon and R5 act independently of each other, allowing helicon to gate locomotion and update the heading direction through excitation of EPG neurons (**Fig. 6**), which are part of a navigational system that allows flies to orientate themselves in their surroundings (Kim et al., 2017; Seelig and Jayaraman, 2013; Turner-Evans et al., 2017). At night, oscillating R5 neurons phase-lock helicon activity in a slow rhythm (**Fig. 4, 5**). Here, our theoretical model recapitulates synchronization of single R5 neurons (Raccuglia et al., 2019) and sets the R5 network as the master oscillator to entrain helicon SWA (**Fig. 5**). The slow process of synchronization fits the homeostatic build-up of sleep need over the time of day (Borbely, 1982) and potentially increases the efficacy to filter out visual stimuli when synchronized by associating helicon. As a consequence of coherent network activity, the functionally antagonistic inputs of helicon and R5 to the head direction system cancel each other out (**Fig. 7**). Indeed, we show that artificially imposing R5 activity at 1 Hz is sufficient to promote quiescence (**Fig. 3**) and impede behavioral responsiveness (**Fig. 5**) to visual stimuli. Moreover, the inhibitory tone on EPG neurons mediated by oscillating R5 neurons at night reduces the functional connectivity of helicon to EPG, which might prevent updating of the heading direction and the initiation of behavioral programs, effectively reducing behavioral responsiveness (**Fig. 7**). While similar interactions have recently been proposed as part of a theoretical model (Flores-Valle et al., 2021), we here provide experimental evidence for this, demonstrating that R5 and helicon antagonistically regulate the head direction system (**Fig. 6**). Salient sensory stimuli might be able to break up synchronized states by engaging additional neurons or specific engrams, thus acutely increasing functional connectivity, which might lead to flies waking up and reacting to external stimuli (French et al., 2021). We speculate that the neural interactions we discovered in this work are likely not restricted to flies, but could represent a neural filtering strategy (Tononi and Massimini, 2008) that correlates to a quiescent state less influenced by external cues, facilitating the shift from an allocentric-like to an egocentric-like worldview.

### Neural networks integrating sleep need, navigation and postural control

One of the main functions of SWA during sleep has been shown to be the promotion of synaptic plasticity to consolidate experiences made during the day, providing the foundation for long-term memories (Klinzing et al., 2019). The circuitry we investigated in this work unifies visual processing, locomotion, navigation and sleep regulation. Ring neurons in general have been shown to play a role in visual place learning (Ofstad et al., 2011) and it is likely, that the oscillatory coherence reported here facilitates synaptic plasticity within the navigational system. Theoretical work has proposed that R5 could balance Hebbian plasticity in the navigational system by alternating between oscillatory and non-oscillatory states and by modulating the synaptic weights to head direction neurons (Fisher et al., 2019; Flores-Valle et al., 2021; Kim et al., 2019; Lu et al., 2022; Lyu et al., 2022; Vafidis et al., 2022).

Besides their role in navigation, we found that frequency-specific activation of R5 to mimic SWA induced a quiescent state accompanied by grooming (**Fig. 3**). Interestingly, we also observed frequent grooming during quiescence that occurred after a period of hyperactivity induced by artificial activation of the head direction system (**Fig. 6**), indicating that quiescence and grooming might be associated with each other. Indeed, grooming behavior in *Drosophila* is regulated by the circadian clock (Qiao et al., 2018) and occurs predominantly during short resting phases prior to night time sleep. Therefore, grooming could be considered as a transitioning stage for the egocentric mode (Qiao et al., 2018) and the activation of R5 might promote this preparatory stage eventually leading to sleep.

While optogenetic activation of R5 at 1 Hz induces quiescence, we found that activating R5 at 10 Hz can lead to a complete loss of postural control with flies falling over (**Fig. 3, 7**). Interestingly, flies also lost postural control when the head direction neurons were continuously hyperpolarized indicating that inhibitory control mediated by R5 SWA inhibits locomotion and provides the neurophysiological basis for maintaining postural control during quiescence (**Fig. 7**). Moreover, SWA provides the temporal structure to create a ‘breakable’ filter, allowing strong or salient stimuli to ‘break’ the neural interaction and allow the animal to wake up. We propose that oscillatory patterns serve as incomplete ‘road blocks’ and allow for shutting off the route to the whole brain while not leading to unresponsiveness as observed in aberrant brain states such as coma. We therefore identified a network motif that not only mediates sleep regulation but also provides a system for the central regulation of postural control and locomotion-based behavioral programs.

## STAR METHODS

### KEY RESOURCES TABLE

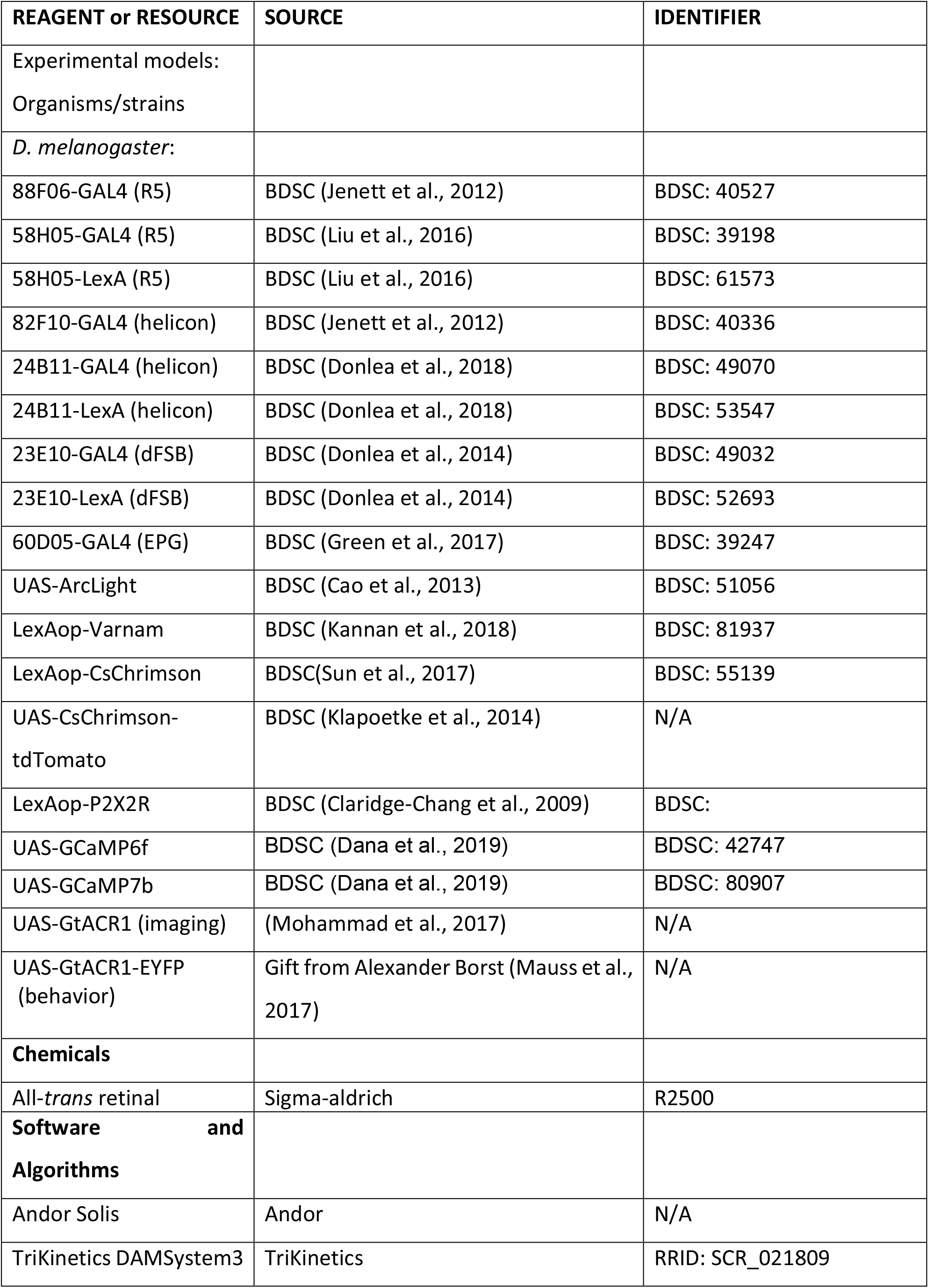
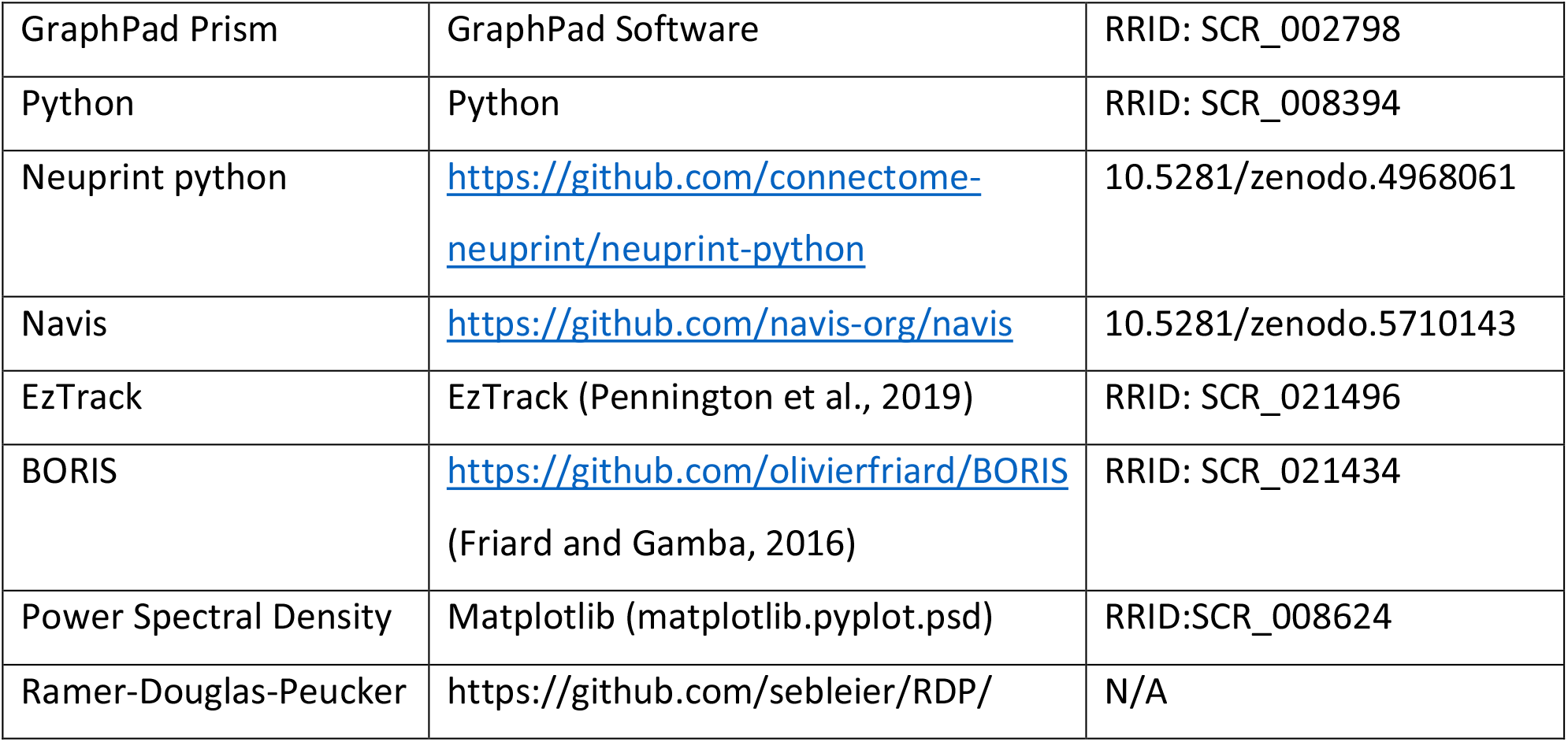

| REAGENT or RESOURCE | SOURCE | IDENTIFIER |
| --- | --- | --- |
| Experimental models:<br>Organisms/strains |  |  |
| *D. melanogaster*: |  |  |
| 88F06-GAL4 (R5) | BDSC (Jenett et al., 2012) | BDSC: 40527 |
| 58H05-GAL4 (R5) | BDSC (Liu et al., 2016) | BDSC: 39198 |
| 58H05-LexA (R5) | BDSC (Liu et al., 2016) | BDSC: 61573 |
| 82F10-GAL4 (helicon) | BDSC (Jenett et al., 2012) | BDSC: 40336 |
| 24B11-GAL4 (helicon) | BDSC (Donlea et al., 2018) | BDSC: 49070 |
| 24B11-LexA (helicon) | BDSC (Donlea et al., 2018) | BDSC: 53547 |
| 23E10-GAL4 (dFSB) | BDSC (Donlea et al., 2014) | BDSC: 49032 |
| 23E10-LexA (dFSB) | BDSC (Donlea et al., 2014) | BDSC: 52693 |
| 60D05-GAL4 (EPG) | BDSC (Green et al., 2017) | BDSC: 39247 |
| UAS-ArcLight | BDSC (Cao et al., 2013) | BDSC: 51056 |
| LexAop-Varnam | BDSC (Kannan et al., 2018) | BDSC: 81937 |
| LexAop-CsChrimson | BDSC(Sun et al., 2017) | BDSC: 55139 |
| UAS-CsChrimson-<br>tdTomato | BDSC (Klapoetke et al., 2014) | N/A |
| LexAop-P2X2R | BDSC (Claridge-Chang et al., 2009) | BDSC: |
| UAS-GCaMP6f | BDSC (Dana et al., 2019) | BDSC: 42747 |
| UAS-GCaMP7b | BDSC (Dana et al., 2019) | BDSC: 80907 |
| UAS-GtACR1 (imaging) | (Mohammad et al., 2017) | N/A |
| UAS-GtACR1-EYFP<br>(behavior) | Gift from Alexander Borst (Mauss et al., 2017) | N/A |
| **Chemicals** |  |  |
| All-*trans* retinal | Sigma-aldrich | R2500 |
| **Software and Algorithms** |  |  |
| Andor Solis | Andor | N/A |
| TriKinetics DAMSystem3 | TriKinetics | RRID: SCR_021809 |
| GraphPad Prism | GraphPad Software | RRID: SCR_002798 |
| Python | Python | RRID: SCR_008394 |
| Neuprint python | https://github.com/connectome-neuprint/neuprint-python | 10.5281/zenodo.4968061 |
| Navis | https://github.com/navis-org/navis | 10.5281/zenodo.5710143 |
| EzTrack | EzTrack (Pennington et al., 2019) | RRID: SCR_021496 |
| BORIS | https://github.com/olivierfriard/BORIS<br>(Friard and Gamba, 2016) | RRID: SCR_021434 |
| Power Spectral Density | Matplotlib (matplotlib.pyplot.psd) | RRID:SCR_008624 |
| Ramer-Douglas-Peucker | https://github.com/sebleier/RDP/ | N/A |

### LEAD CONTACT AND MATERIALS AVAILABILITY

Requests for raw data, instrumentation and analysis code should be directed to the Lead Contact, David Owald. This study did not generate new unique reagents.

### EXPERIMENTAL MODEL AND SUBJECT DETAILS

Flies (*Drosophila melanogaster*) were reared on standard cornmeal food at 25 °C and 60% humidity under a 12 h light/dark regime. Details of genotype and source of flies are given in **Key Resources Table**.

### METHOD DETAILS

#### Imaging of neural activity

For voltage and Ca^2+^ imaging experiments we used 3-10 day old female flies and performed whole-brain explant dissections as previously described (Cao et al., 2013; Raccuglia et al., 2019). If not indicated otherwise, morning experiments were performed at ZT (Zeitgeber time) 2-4 (during a 12/12 h light/dark cycle the onset of light is at ZT 0 h and the offset of light is at ZT 12 h) and night experiments were performed at ZT 14-18.

To record spontaneous activity (**Fig. 1, 2G-I, 4, 6D**), we used a low Mg^2+^ external solution (5 mM) consisting of (in mM) 90 NaCl, 3 KCl, 1.5 CaCl_2_, 5 MgCl_2_, 1 NaH_2_PO_4_, 10 glucose, 10 sucrose, 8 trehalose, 5 TES and 26 NaHCO_3_. To suppress spontaneous activity (**Fig 2A-F, 6G**) we used a high Mg^2+^ external solution (20 mM) consisting of (in mM): 70 NaCl, 3 KCl, 1.5 CaCl_2_, 20 MgCl_2_, 1 NaH_2_PO_4_, 10 glucose, 10 sucrose, 8 trehalose, 5 TES and 26 NaHCO_3_. To compensate for Cl^-^ we adjusted NaCl concentration in the external solution, which did not alter spontaneous activity (data not shown). External solution was adjusted to a pH of 7.4, with an osmolarity of 280 mmol/kg.

Imaging was performed using an Olympus BX51WI microscope using a Plan Apochromat 40×, numerical aperture 0.8, water-immersion objective (Olympus, Japan). The objective C-mount image was projected onto an Andor iXon-888 camera controlled by Andor Solis software. Voltage imaging was performed at ∼80 frames/s while Ca^2+^ imaging was performed at ∼20 frames/s. ArcLight and GCaMP were excited at 475/35 nm using a Lumencor Spectra X-Light engine (Lumencor, USA) LED system. For Ca^2+^ imaging we used either GCaMP6f or GCaMP7b. The increased brightness and sensitivity of GCaMP7b allows for an improved detection of anatomical structures and neural activity in promotor lines with low baseline expression (Dana et al., 2019). To avoid saturated fluorescent images, we adjusted the LED power for each recording individually. To determine absolute fluorescence in single helicon cell bodies (**Fig. S5-1D**) LED power was kept constant at 24% throughout all recordings. For dual color voltage imaging, ArcLight and Varnam were simultaneously excited at 475/35 nm and 542/33 nm using a Spectra X-Light engine LED (Lumencor, USA) system. For separating the emission wavelengths, we used the W-View Gemini image splitter (Hamamatsu, Japan) with an adequate dichroic mirror for ArcLight (520/35 nm) and Varnam (641/75 nm).

For dual-color voltage imaging and for optogenetic experiments, all*-trans* retinal (Sigma-Aldrich) was added to the fly food as a 50 mM stock dissolved in 95 % ethanol. Female flies were collected after eclosion and transferred to fly food containing 1.3 mM of all-trans retinal for 2-4 days. CsChrimson was excited at 637/12 nm at a power density of 190-740 µW/cm^2^. For optogenetic hyperpolarization GtACR1 was excited at 542/33 nm at a similar power density. Due to potential spectral overlap, excitation of ArcLight and GCaMP at 475/35 nm was kept minimal (20-170 µW/cm^2^). To reduce optical artefacts generated by red light stimulation we used a custom-made dualband beamsplitter filter cube allowing red light to pass the excitation filter but not the emission filter. To further reduce optical artifacts we performed a background subtraction on imaging recordings (see quantification and statistical analysis).

#### Single-fly tracking experiments

For measuring locomotor activity, we used female flies that were collected within 2-3 days after eclosion. Experiments were performed in the dark in circular arenas (2.5 cm diameter) illuminated from below using a low power infrared (850 nm) LED panel (2.7 mW/cm^2^ at 520 nm). Flies were recorded from above at 15 frames/s using the high-resolution camera Flea3 USB3 (FL3-U3-13E4C-C, Teledyne FLIR) attached to a 25 mm focal length lens (#63-246, Edmund Optics) and an infrared filter (10000314, Neewer) as done elsewhere (Aso and Rubin, 2016). For optogenetic experiments (**Fig. 3, 5** and **6**), flies were reared for 2-4 days after collection in food supplemented with 1.3 mM all-*trans* retinal. To avoid optogenetic activation during this time, vials containing flies were partially covered with a foil which still allowed them to perceive background light. All experiments were carried out at room temperature and during the subjective day of the fly (from ZT1-10).

At the start of each experiment, flies were allowed to acclimatize for 10 min. We then measured the flies’ initial baseline locomotor activity for 5 min (**Fig. 3B**). For optogenetic activation experiments, flies were then exposed to 5 min of 5-ms pulses of red light (∼20 mW/cm^2^ at 632 nm) from below at a specific frequency indicated in the figure legends. Subsequently locomotor activity was recorded for another 10 min. For optogenetic hyperpolarization, flies were exposed to constant 1-min long green light from below (∼9mW/ cm^2^ at 520nm). For measuring absolute locomotor activity (distance travelled) and mean velocities, we considered all movements. To determine whether flies were perturbed in their ability to walk, we only considered velocities equal or higher than 0.25 cm/s (walking velocity and % of time walking) (Donlea et al., 2018; Kottler et al., 2019). In order to normalize locomotor variables to the baseline activity we computed the ‘effect’ of the optogenetic protocol on baseline locomotion [ie. effect on total distance = (distance travelled during stimulation – distance travelled during baseline) / (distance travelled during stimulation + distance travelled during baseline)]. To evaluate turning behavior (**Fig. 6**), we simplified flies’ trajectories using an implementation of the Ramer-Douglas-Peucker algorithm (https://github.com/sebleier/RDP/), computed the direction vectors of the simplified trajectories and classified changes in direction equal or higher than 47° as turns. For high resolution analysis of behavior during R5 activation and EPG inhibition experiments (**Fig. S3F, G and S6D, E**), we manually annotated the occurrence of different behaviors: walking, grooming, resting (flies are immobile with an upright body posture), loss of posture (flies are immobile and upright body posture is lost) and recovering (flies are moving and trying to recover upright body posture).

#### Behavioral responsiveness

To test visual responsiveness of sleep deprived flies (**Fig. 5F-H and S5I, J**), vials containing flies reared in retinal food for 2-3 days were fixed to an analog Multi-Tube Vortexer (VX-2500, VWR) controlled by TriKinetics acquisition software. For sleep deprivation a mechanical stimulus lasting 2 s was provided randomly within a time window of 20 s from the beginning of the subjective night (ZT 20) to the start of the experiment (ZT 1-7). Only sleep deprived flies that did not move in the last 5 min of recording baseline activity (meeting the generally accepted criteria for sleep) were included in the analysis. To assess visual responsiveness during optogenetic activation flies were exposed to a 1-min green light stimulus from above (0.37 mW/cm^2^ measured at 520 nm). Responsiveness to light was evaluated by measuring changes in a fly’s velocity (‘effect on mean velocity’). To determine whether flies were awakened by light we only considered movements equal or higher than 0.25 cm/s.

To test responsiveness to an air puff during optogenetic activation (**Fig. S3D**), rested flies were individually loaded in Trikinetics glass tubes (5×65 mm) connected to an air pump on one side and sealed with perforated parafilm on the other end to allow air flow. Tubes were attached to the LED panel for 1 Hz optogenetic activation and exposed to a constant 10 s air puff (∼10 l/min) that was sufficiently strong to startle the flies. Responsiveness was measured by comparing changes in mean velocity before and after the air puff (40 s each). To exclude movements during the air puff, data 5 s before and after stimulus was not included in the analysis.

#### Network simulations

To investigate the network configurations and mechanisms generating slow-wave oscillations and networks coherence within and between R5 and helicon, we simulated network interactions using a spiking neuron model introduced by Izhikevich (Izhikevich, 2003) (**Fig. 5** and **Fig. S5-1**). For detailed description of the computational model, please see **Supplemental Experimental Procedures**.

#### Connectome analysis

Neuronal types can be found in neuprint (neuprint.org) under the same name used in the figures, with the exception of R neurons (ER5, ER4, etc. in the connectome) and helicon cells (ExR1). The neuron type EPGt (Hulse et al., 2021) was not included in the EPG analysis. In the connectivity plots, input weight contribution of a neuron (or set of neurons) *A* from a neuron (or set of neurons) *B* refers to the number of synaptic connections that *A* receives from *B*, normalized to the total number of input connections of *A*. Output weight contribution of *A* to *B* refers to the number of output synaptic connections from *A* to *B*, normalized by the total number of output connections of *A*. The 2D morphological renderings of neurons were visualized using python. For the analysis of neighboring synapses in EPG neurons, we located individual R5-to-EPG and helicon-to-EPG input connections in every EPG. Then, we identified the closest input connection to each R5-to-EPG and helicon-to-EPG connection in the same neuron (excluding other connections from R5 and helicon cells, respectively and connections from cells not typed in the connectome). Distances between synaptic connections were calculated using geodesic (along the arbor) distance.

#### Visualization, software and statistics

Activity recordings were analyzed using NOSA, a custom-made software specifically developed to analyze voltage and Ca^2+^ imaging recordings (see https://github.com/DavideR2020/NOSA). Relative fluorescence, background subtraction, power spectrum analysis and cross correlation analysis were performed with algorithms described in detail previously (Oltmanns et al., 2020; Raccuglia et al., 2019). From NOSA, data sets were exported to Microsoft Excel for further analysis. For visualization of experimental data, we used CorelDRAW2020. For a more intuitive display, all ArcLight and Varnam recordings are inverted, as depolarization leads to a decrease in fluorescence intensity while hyperpolarization leads to an increase. For power spectrum analysis, the unit of the power spectral density (PSD) is indicated as PSD = [(ΔF/F)^2^ /Hz] *100. To quantify response amplitudes during optogenetic activation (**Fig. 6H**) we computed the difference of the average relative change in fluorescence intensity during and before optogenetic stimulation. All imaging data are presented as means with standard error of the mean (SEM).

For tracking recordings, single fly position data was extracted from videos using EzTrack (Pennington et al., 2019) (https://github.com/denisecailab/ezTrack). Hemibrain connectome data (version 2.1) (Scheffer et al., 2020) was accessed using neuprint-python (https://github.com/connectome-neuprint/neuprint-python) and geodesic (along the arbor) distances between synaptic connections were calculated using navis (https://github.com/navis-org/navis). Manual annotation of behaviors was performed using BORIS (https://github.com/olivierfriard/BORIS) (Friard and Gamba, 2016). Behavioral and connectome data were analyzed using custom-made python scripts. In behavioral panels, box-plots show median (line), quartiles (boxes) and range (whiskers) and locomotor activity curves indicate mean ± SEM. Behavioral and connectome data were visualized using python.

R5 and helicon network simulations were performed, analysed and visualized using python.

Statistical analysis was carried out in GraphPad Prism. In all panels, significance is represented as follows: *p ≤ 0.05, **p ≤ 0.01, ***p ≤ 0.001 and ns (not significant). The statistical tests performed in each panel are indicated in the figure legends. For unpaired comparisons, we used Mann-Whitney test when comparing 2 groups and Kruskal-Wallis test with Dunn’s post hoc analysis for multiple comparisons. For paired comparisons, we used Wilxocon matched pairs signed rank test when comparing 2 groups and Friedman test with Dunn’s post hoc analysis for multiple comparisons. Comparisons to 0 were performed using Wilcoxon signed-ranked tests and percentages were compared using Binomial test.

## DATA AND CODE AVAILABILITY

Raw data are available from the corresponding author upon reasonable request.

## Acknowledgements

We thank Peter Spende, Alexej Schatz, Waldemar Krzok, Raz Flieshman, Karim Izzuldin and Katharina Stumpenhorst for help with the behavioral arena; Johannes Felsenberg and members of the Owald lab for comments on the manuscript, Scidraw for allowing us to modify and use their schematics (fly drawing: doi.org/10.5281/zenodo.3926137) and Alexander Borst, the Janelia fly line project and Bloomington Stock Center for flies.

## Author contribution

Conceptualization: D.R., R.S-G., L.K., R.K. and D.O.

Investigation: D.R., R.S-G., L.K, A.E., C.B., S.R.J, and G. Y-D.

Resources: D.R., Y.W, R.K., J.R.G. and D.O.

Writing – Original Draft: D.R., R.S-G., L.K., R.K. and D.O.

Writing – Review & Editing: A.E, C.B. and J.R.G.

Authors declare no competing interests.

---

**Figure S1.**
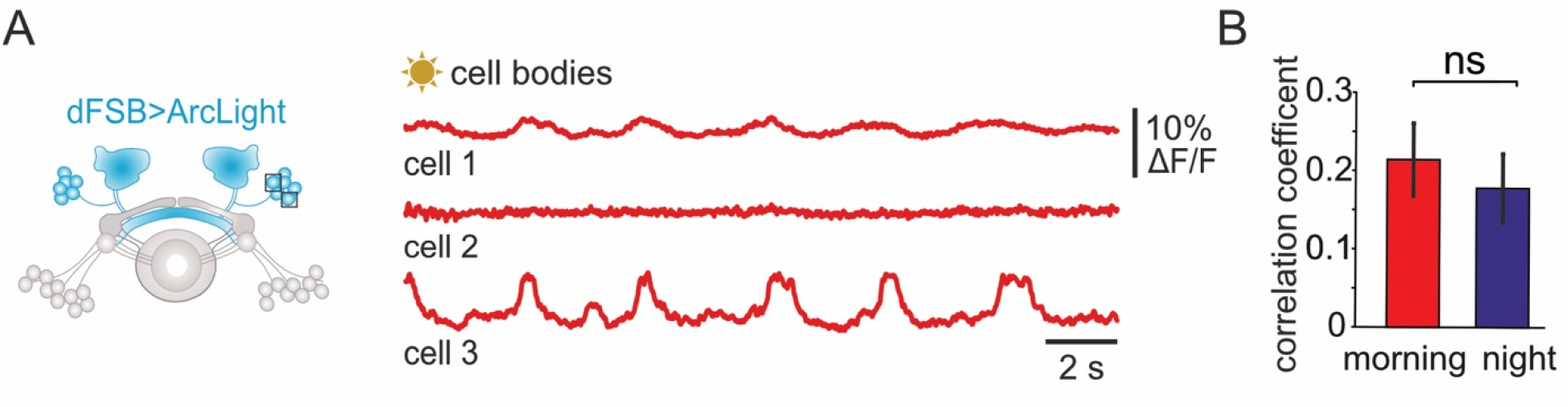
**(A)** *Ex vivo* multi-cellular voltage imaging of individual dFSB neurons recorded in the morning (ZT 2-4). Squares schematically indicate recording sites. **(B)** Correlation between electrical patterns of individual cell bodies is similar in the morning and at night (*n*=30-39, Mann-Whitney test).

**Figure S2.**
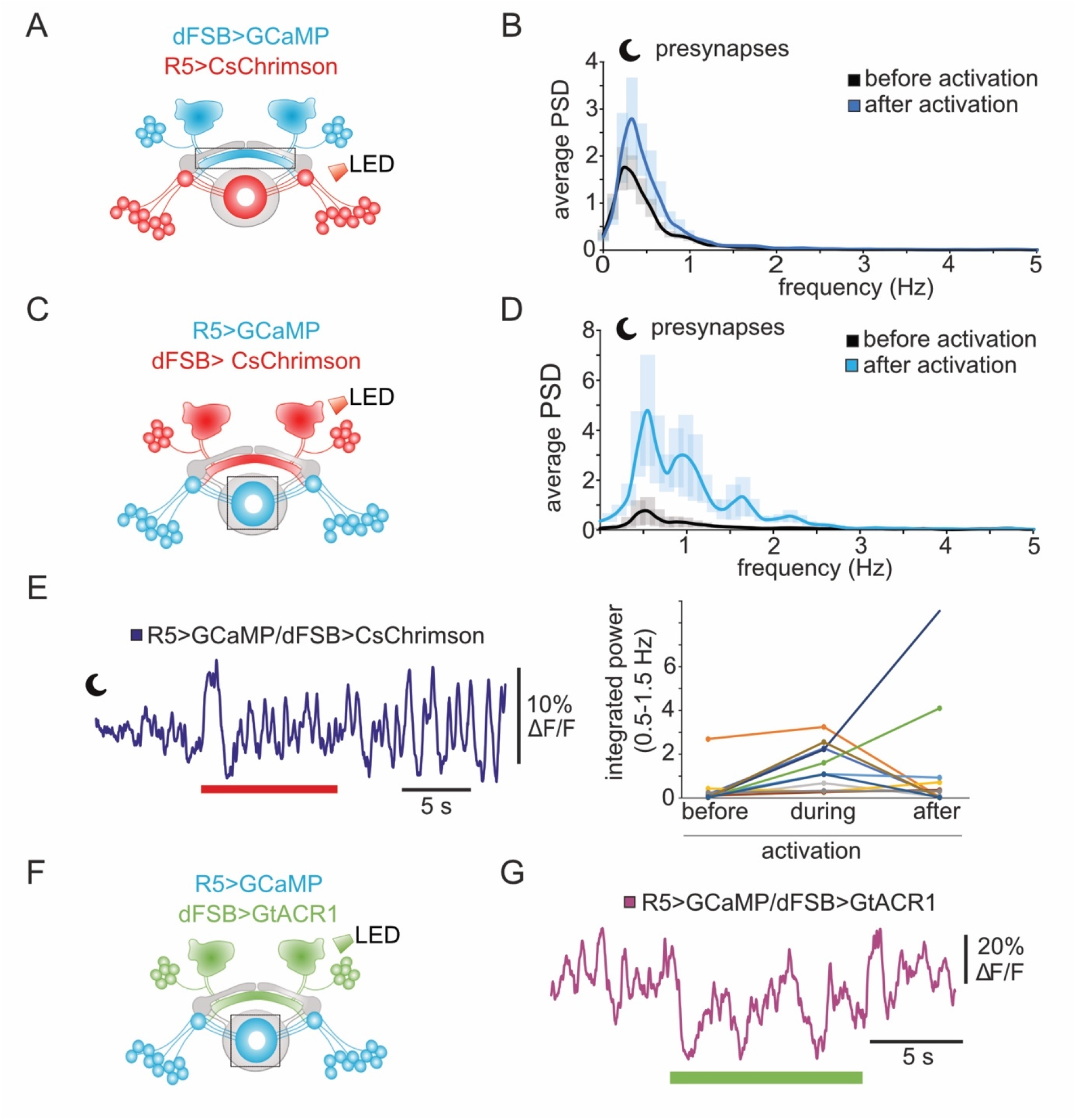
**(A, C)** Schematic overview of presynaptic recording sites and sites of optogenetic activation. **(B)** Average power spectra of dFSB compound oscillations before and after optogenetic stimulation of R5 (*n*=10). **(D)** Average power spectra of R5 compound oscillations before and during optogenetic stimulation of dFSB neurons (*n*=11). **(E)** Left: example recording of ongoing R5 presynaptic Ca^2+^ oscillations after cessation of dFSB stimulation. Right: individual data points of R5 slow-wave power (0.5-1.5 Hz) show that in some cases R5 oscillations persist after cessation of dFSB stimulation. **(F)** Schematic overview indicating expression of GtACR1 in dFSB and Ca^2+^ recording sites at the R5 ring. **(G)** Example recording of R5 presynaptic Ca^2+^ activity showing partially recovering oscillatory activity during optogenetic inhibition of dFSB neurons.

**Figure S3.**
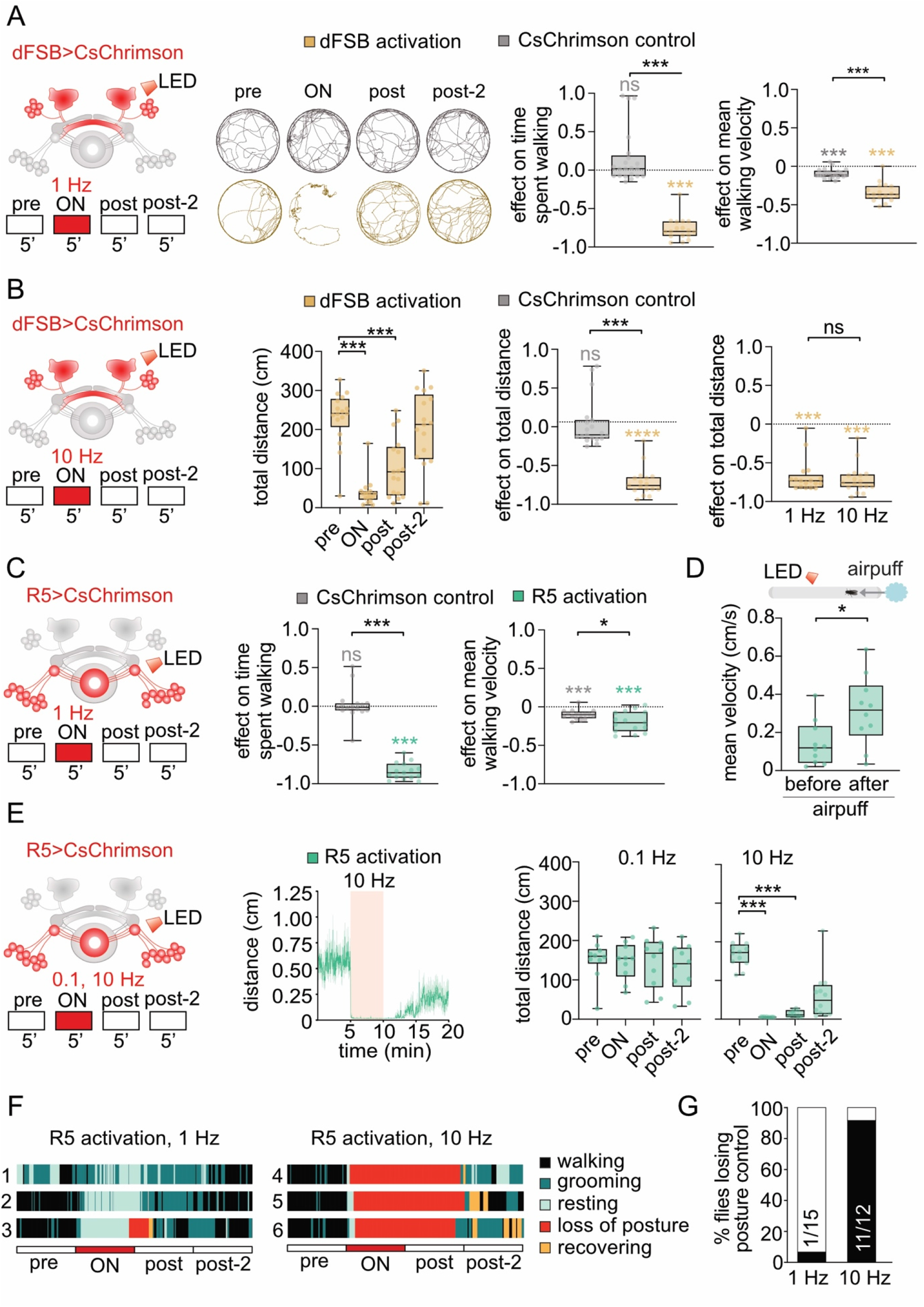
**(A)** Optogenetic stimulation of dFSB neurons at 1 Hz using the protocol described in Fig. 3C. Example trajectories of CsChrimson control (+ retinal) and 23E10-GAL4>UAS-CsChrimson (+ retinal) flies. Optogenetic stimulation of dFSB neurons at 1 Hz in rested flies decreased percentage of time walking and mean walking velocity compared to control flies, *n*=17-19 each group, Mann-Whitney test (comparisons between genotypes depicted in black) and Wilcoxon signed-rank test (comparisons to baseline locomotion depicted in color). **(B)** Activation of dFSB at 10 Hz decreased total distance travelled during and after activation (*n*=17, Friedman test with Dunn’s multiple comparisons). The effect of dFSB stimulation at 10 Hz on distance travelled differed to that of control flies, *n*=17-19 each group, Mann-Whitney test (comparisons between genotypes depicted in black) and Wilcoxon signed-rank test (comparisons to baseline locomotion depicted in color). The effect of dFSB stimulation on distance travelled did not differ between different frequencies of activation (*n*= 17 each, Mann-Whitney test). **(C)** Activation of R5 neurons at 1 Hz in rested flies expressing CsChrimson under 58H05-GAL4 control (+ retinal) decreased percentage of time walking and mean walking velocity compared to control flies, *n*=15-16 each group, Mann-Whitney test (comparisons between genotypes depicted in black) and Wilcoxon signed-rank test (comparisons to baseline locomotion depicted in color). **(D)** Exposure to a 10 s-long air puff during R5 activation at 1 Hz increased the mean velocity of rested flies in 5×65 mm tubes (+ retinal, *n*=10, Wilcoxon matched-pairs signed rank test). **(E)** Average locomotion during activation of R5 neurons at different frequencies. Curve: mean ± SEM. Activation of R5 neurons in rested flies at 0.1 Hz did not affect locomotion, whereas 10 Hz stimulation reduced distance travelled during and after activation (*n*=9-12 each group, Friedman tests with Dunn’s multiple comparisons). **(F)** Raster plots of six example flies (58H05-GAL4>UAS-CsChrimson) before, during and after R5 activation at 1 Hz (left) or 10 Hz (right). Colors indicate five designated behaviors: walking, grooming, resting (flies are immobile), loss of posture (flies are immobile and upright body posture is lost) and recovering (flies are recovering locomotion and upright body posture). **(G)** Fraction of flies that lost postural control during R5 activation at different frequencies (black). The *n* of flies for each condition is indicated in the bar plots.

**Figure S4.**
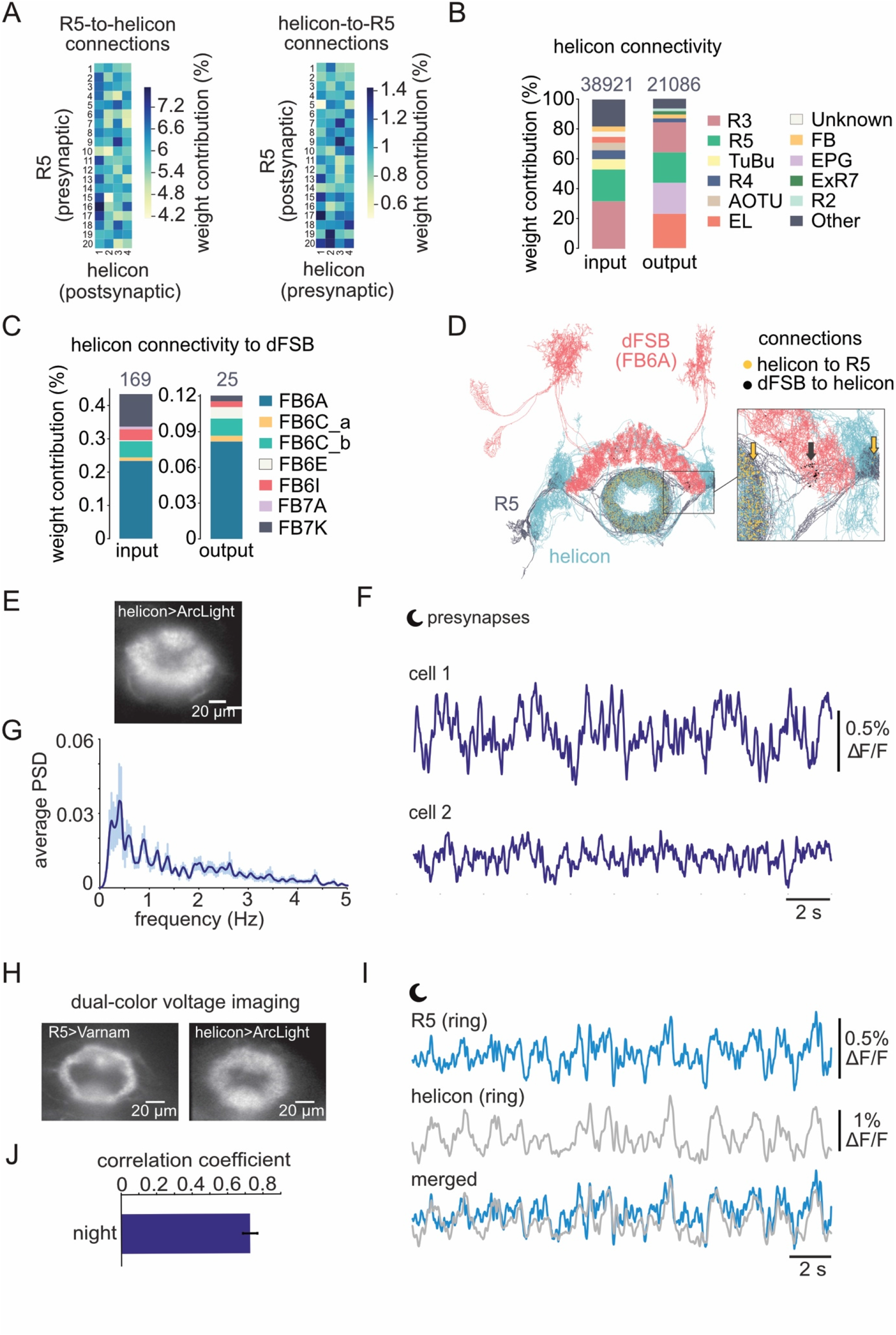
**(A)** Neuron-to-neuron connectivity plot for connections from R5 to helicon (left) and vice versa (right). Number of connections were normalized to the total number of output connections of each R5 neuron (left) and to the total number of output connections of each helicon cell (right). **(B)** Top synaptic input and output connections of all helicon cells. Number of input and output connections of helicon per neuronal type were normalized to the total number of input and output connections of helicon cells. **(C)** Fraction of helicon cell input and output from dFSB neuron types normalized to the total number of input and output connections of helicon cells. **(D)** Morphological rendering of R5 neurons, helicon cell and FB6A (dFSB neuron type) neurons. Yellow dots/arrow indicate areas of synaptic connections of helicon cells to R5 neurons and black dots/arrow indicate areas of synaptic connections from the dFSB to helicon. **(E)** Wide-field image of presynapses expressing the green voltage indicator ArcLight in helicon cells (24B11-GAL4). **(F)** Example recordings of helicon presynaptic electrical patterns recorded at night (ZT 14-16). **(G)** Average power spectrum of electrical compound presynaptic activity recorded at night (n=6, ZT 14-16). **(H)** Wide-field images of presynaptic structures expressing the green voltage indicator ArcLight in helicon cells (24B11-GAL4) and the red voltage indicator Varnam in R5 neurons (58H05-LexA). **(I)** Example recordings of R5 and helicon presynaptic electrical patterns recorded at night (ZT 14-16). **(J)** Correlation between electrical patterns of helicon and R5 at night (n=8).

**Figure S5-1.**
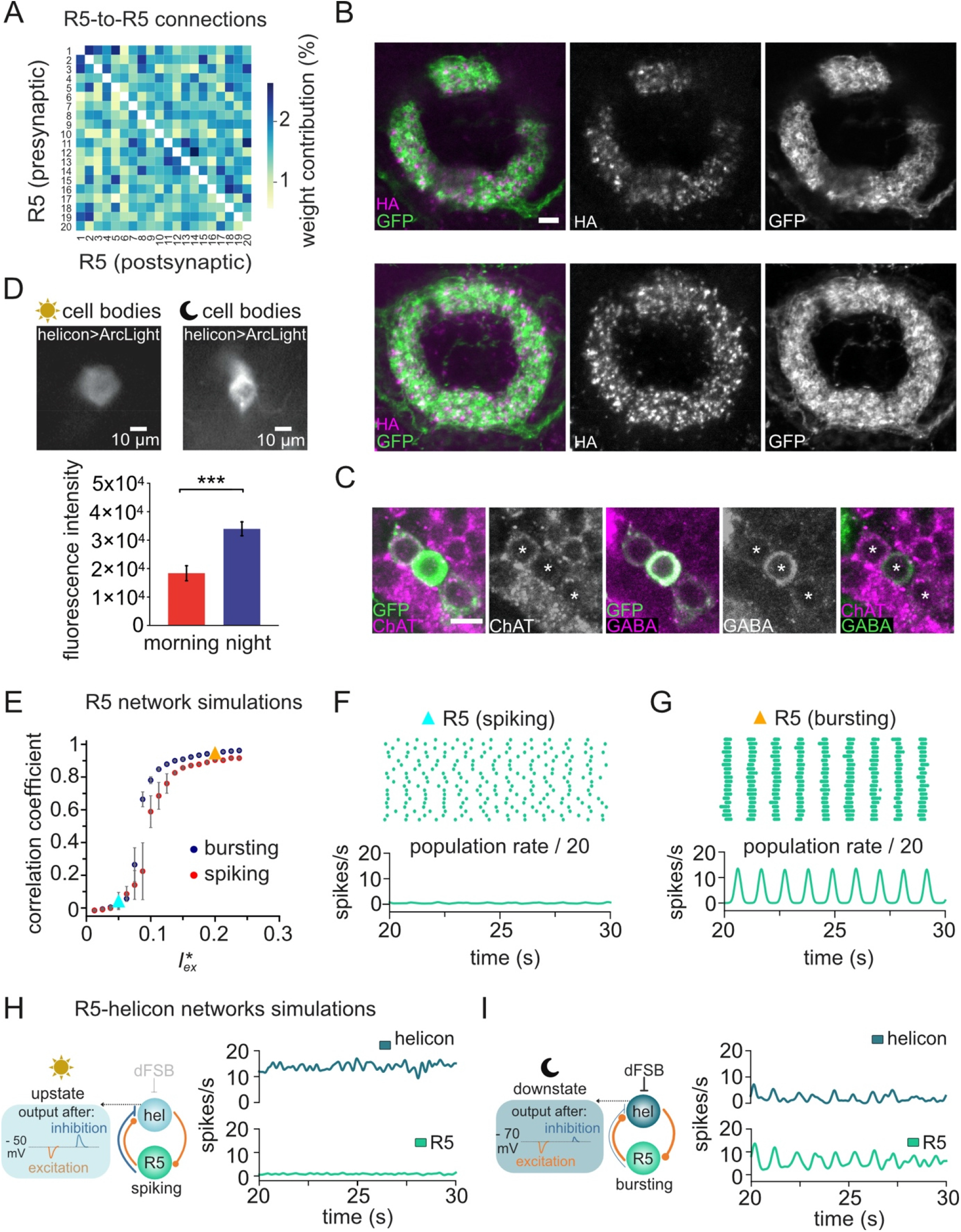
**(A)** Neuron-to-neuron connectivity plot for connections from R5 to R5. Number of connections were normalized to the total number of output connections of each R5 neuron. White squares indicate no direct connections. **(B)** Upper row: expression of vACht::HA and mCD8::GFP in presynaptic sites of R5 neurons targeted by 58H05-GAL4 and immunostained with anti-GFP and anti-HA. Single confocal plane. Bottom row: Maximum projection of R5 presynaptic region. Scale bar: 5 μm. **(C)** Single confocal sections of R5 cell bodies expressing mCD8::GFP under control of 58H05-GAL4 labeled with antibodies against ChAT and GABA. Images are representative examples of one brain from four independent experiments. Asterisks indicate GFP expressing R5 neurons. Scale bar: 5 μm. **(D)** Top: wide-field images of single helicon cell bodies expressing ArcLight (24B11-Gal4) in the morning (ZT1-4) and at night (ZT12-16). Average fluorescence intensity is increased at night, indicating that helicon cells are more hyperpolarized at night relative to the morning (*n*=12, Mann-Whitney test). **(E)-(I)** Numerical simulations of R5 and helicon networks; for details on the implementation, see Supplemental Experimental Procedures. **(E)** Simulated synchronization in an isolated R5 network model. Mean correlation coefficients of 20 recurrently connected R5 neurons (spiking or bursting) for a fixed inhibitory coupling strength (−0.01) and varying excitatory coupling strength *h^∗^*. Network synchronization correlated with strength of excitation, suggesting that the interaction of R5 neurons has some excitatory component and/or that other connecting networks provide excitation. Colored triangles indicate the mean correlation coefficients used in **F** and **G** (n=10 simulations each). **(F)** Top: spike raster plot of 20 R5 neurons in spiking mode and in a rather desynchronized state (*h^∗^* = 0.05, mean correlation coefficient of ∼ 0.3). Bottom: population rate divided by the number of neurons (n=10 simulations). **(G)** Top: spike raster plot of 20 R5 neurons in burst-firing mode and in a highly synchronized state (*h^∗^* = 0.20, mean correlation coefficient of ∼0.9). Bottom: population rate divided by the number of neurons (n=10 simulations). **(H)** Left: schematic summary of simulated R5 and helicon interactions in the morning (see Fig. 5A). Right: compound activity of helicon and R5 neurons in the morning normalized by the number of neurons in each network. In the morning, SWA was weak in both networks (n=10 simulations). **(I)** Left: schematic summary of simulated R5 and helicon interactions at night (see Fig. 5A). Right: compound activity of membrane potential dynamics of helicon cells and R5 neurons at night (see Fig. 5A) normalized by the number of neurons in each network. In the evening, both networks showed increased SWA activity (n=10 simulations).

**Figure S5-2.**
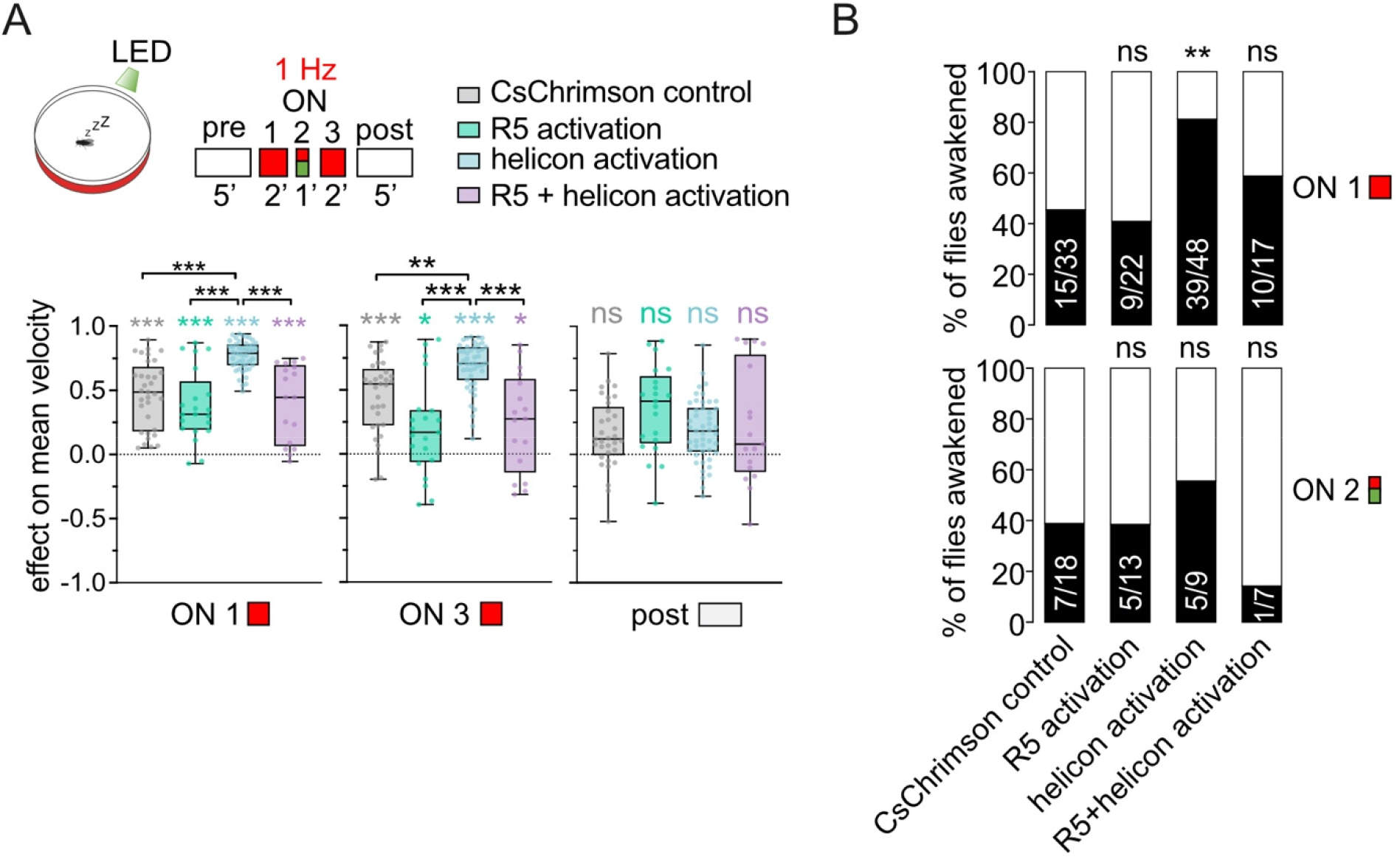
**(A)** Top: schematic overview of optogenetic activation protocol to test behavioral responsiveness during network activation in sleep deprived flies (described in Fig. 5F). Bottom: effect of optogenetic stimulation at different times of the experiment. The effect of helicon activation on locomotion differed to that of R5 and R5+helicon activation, *n=33* for CsChrimson control, *n*=22 for 58H05-LexA>LexAop-CsChrimson, *n*=47 for 24B11-LexA>LexAop-CsChrimson and *n*=17 for 58H05-LexA;24B11-LexA>LexAop-CsChrimson, Kruskal-Wallis test with Dunn’s multiple comparisons (comparisons between genotypes, black asterisks) and Wilcoxon signed-rank tests (comparisons to baseline locomotion, colored asterisks). Behavioral data was acquired during the subjective day of the flies (ZT 1-7) after 12-19 h of sleep deprivation. **(B)** Percentage of sleeping flies that started walking (‘awakened’) at the beginning of the optogenetic stimulation (top, ‘ON 1’) or when exposed to green light (bottom, ‘ON 2’). 1 Hz stimulation of the helicon network led to more flies waking up at the beginning of the stimulation when compared to the genetic control, Binomial test. The *n* of flies for each condition is indicated in the bar plots.

**Figure S6.**
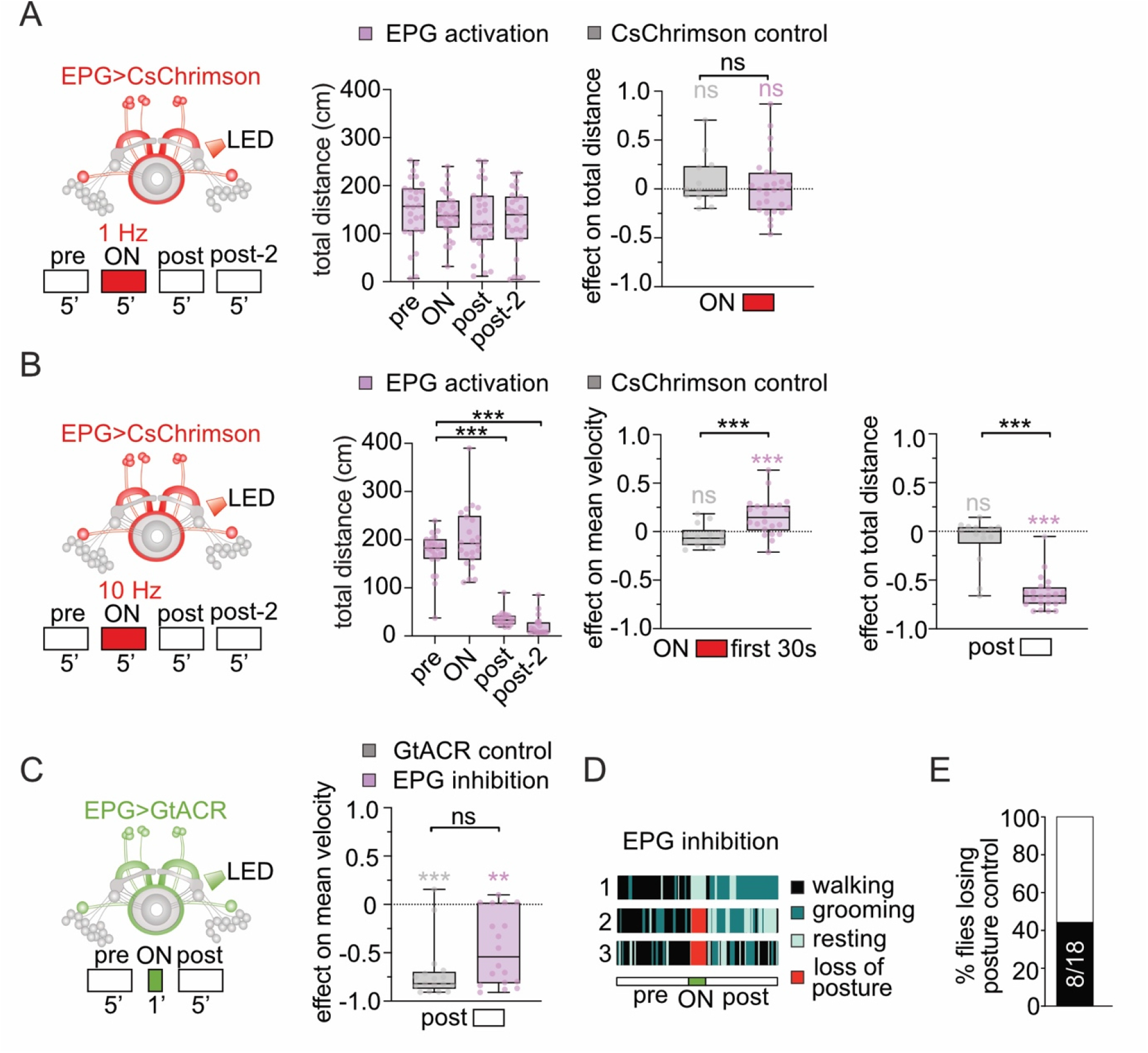
**(A)** Schematic overview of the optogenetic protocol used to activate EPG neurons at 1 Hz for the 60D05-GAL4 control using the stimulation protocol described in Fig. 3C. Activation of EPG neurons in rested flies at 1 Hz did not affect locomotion (*n*=27, Friedman test with Dunn’s multiple comparisons). The effect on distance travelled of 1 Hz stimulation in 60D05-GAL4>UAS-CsChrimson flies (+ retinal) did not differ from that of control flies (+ retinal), *n*=13-27 each group, Mann-Whitney test (comparisons between genotypes depicted in black) and Wilcoxon signed-rank test (comparisons to baseline locomotion depicted in color). **(B)** Schematic overview of 10 Hz EPG activation under 60D05-GAL4 control in rested flies. High frequency activation of EPG neurons reduced locomotion after stimulation, (*n*=22, Friedman test with Dunn’s multiple comparisons). 10 Hz EPG activation led to an increased mean velocity in the first seconds of stimulation and decreased distance travelled after stimulation compared to control flies, *n*=14-22 each group, Mann-Whitney test (comparisons between genotypes depicted in black) and Wilcoxon signed-rank test (comparisons to baseline locomotion depicted in color). **(C)** Schematic of constant inactivation of EPG neurons expressing UAS-GtACR1. EPG block did not affect mean velocity after optogenetic inhibition, *n*=17 for GtACR1 control and *n*=19 for EPG inhibition, Mann-Whitney test (comparisons between genotypes depited in black) and Wilcoxon signed-rank test (comparisons to baseline locomotion depicted in color). **(D)** Raster plots of three example flies (60D05-GAL4>UAS-CsChrimson) before, during and after EPG inhibition. Colors indicate four designated behaviors: walking, grooming, resting (flies are immobile) and loss of posture (flies are immobile and upright body posture is lost). **(G)** Fraction of 60D05-GAL4>UAS-CsChrimson flies that lost postural control during EPG inhibition (black). The *n* of flies is indicated in the bar plot.

