## Supplementary information for "Coherent multi-level network oscillations create neural filters to favor quiescence over navigation in *Drosophila*"

### SUPPLEMENTAL EXPERIMENTAL PROCEDURES

#### Network simulations (referring to Fig. 5A-E and S5-1E-I)

*Neuron model*

To keep models of spiking networks as simple as possible, neurons are described by the framework introduced by Izhikevich (Izhikevich, 2003), which can simulate a wide variety of spiking behaviors while being much simpler than conductance-based models. The Izhikevich model consists of the two coupled differential equations

$$\frac{dv}{dt} = [0.04v^2 + 5v + 140] \cdot 0.05 - u + I + I_{syn}$$

$$\frac{du}{dt} = [a(bv - u)] \cdot 0.05$$

with parameters described in **Table 1**. Every time a neuron's membrane potential $v$ reaches a threshold $v_{th}$ a spike is emitted, and $v$ is set to the reset potential $c$. Moreover, the recovery variable $u$, which accounts for or the activation of $K^+$ and inactivation of $Na^+$ currents, is increased by the amount $d$. The variables $I$ and $I_{syn}$ denote intrinsic and synaptic currents, respectively. The intrinsic current $I$ influences the firing rate of a neuron; to introduce variability of spiking, after each spike we reset $I$ to a new constant value $I = I_0 + \sigma \cdot N(0,1)$ where $I_0$ denotes the mean current, $N(0,1)$ is a standard normal distribution, and the parameter $\sigma$ scales the level of noise, with regular (oscillatory) activity for low noise (typical for R5) (Liu et al., 2016) and irregular activity for high noise (typical for helicon) (Donlea et al., 2018). Activity could arise as single spikes (typical for R5 during day) or bursts (typical for R5 during night) (Liu et al., 2016), which depends also on other model parameters (mainly $d$). To achieve slow-wave-like oscillation frequencies in the range of ~1 Hz and spike waveforms resembling those of neurons in *Drosophila* (Raccuglia et al., 2019), we slowed down the timescale of the original Izhikevich model (designed for cortical neurons) by factors 0.05 in the two equations. Note that all dynamic variables and parameters of the model are unitless; nevertheless, numerical values are chosen such that voltage $v$ can be interpreted in millivolts and time $t$ in milliseconds. In the neuron model described in the two differential equations, $a$ denotes the time-scale of the recovery variable, $b$ is the sensitivity of the recovery variable to sub-threshold fluctuations of the membrane potential, $c$ is the after spike reset value of the membrane potential, $d$ represents the effect of slow $K^+$ and $Na^+$ conductances on the membrane potential recovery post-spike, $I_0$ denotes the mean of the intrinsic current, and $\sigma$ scales the level of noise. The voltage threshold is $v_{th}$.

| | $a$ | $b$ | $c$ | $d$ | $I_0$ | $\sigma$ | $v_{th}$ |
|---|---|---|---|---|---|---|---|
| **R5 during day** | 0.02 | 0.2 | −65 | 6 | 0.34 | 0.02 | −10 |
| **R5 at night** | 0.02 | 0.3 | −50 | 1.6 | 0.3 | 0.08 | −10 |
| **helicon during day** | 0.02 | 0.2 | −65 | 6 | 4.5 | 5 | −10 |
| **helicon at night** | 0.02 | 0.2 | −65 | 6 | −0.75 | 5 | −10 |

**Table 1.** Parameters of R5 and helicon model neurons during day and night.

*Synapse model*

Interactions between neurons are described by postsynaptic currents that are summarized by $I_{syn}$. Each time $t_s$ a presynaptic neuron fires, we add to $I_{syn}$ a transient current $I^*[(t-t_s)/\tau]\exp[-(t-t_s)/\tau]$ which is an alpha function with time constant $\tau$ and amplitude $I^*$. The total charge transfer (i.e. the integral over time) is $I^*\tau$ . The sign of $I^*$ depends on whether the input is excitatory ($I^*_{exc} > 0$) or inhibitory ($I^*_{inh} < 0$). Excitatory postsynaptic currents are modeled with a faster time constant $\tau_{exc} = 4$ ms (based on patch-clamp recordings of R5, data non shown), whereas inhibitory postsynaptic currents are slower with time constants $\tau_{inh} = 20$ ms (Lee et al., 2003).

*Network model and connectivity*

As network simulations of a purely inhibitory R5 network showed no synchronization (example in **Fig. S5-1E**) and experimental data indicated an excitatory output component of R5 (**Fig. S5-1B, C**), we assumed a network of 20 R5 neurons that are equally excitatory and inhibitory. Thus, the postsynaptic currents elicited by spiking of a single R5 neuron are described by the superposition of a faster excitatory current and a slower inhibitory current. The network of 4 helicon cells was assumed to be excitatory and firing was tuned to be irregular. The number of neurons in each network and the recurrent connectivity within and between networks (all-to-all in all cases) were based on connectome data (Hulse et al., 2021). Note that the magnitudes of connection strengths *I** in the modelled network were scaled by the number of synaptic connections in the connectome, which added some variability.

*Coupling at day and night*

We considered a morning and night configuration to account for different R5 and helicon network activity states. In the morning configuration, R5 neurons are in spiking mode (Liu et al., 2016) and helicon cells are in a more active state (‘upstate’) characterized by higher firing rates (~17 spikes/s) and a more depolarized membrane — likely due to a lower inhibition from the dFSB (Donlea et al., 2018). In this upstate, we assume that helicon cells’ resting membrane potentials are way above the chloride reversal potential, which renders them responsive to both inhibitory and excitatory input from R5. In the night configuration, R5 neurons are in a burst-firing mode (Liu et al., 2016) and helicon cells are in a less active state (‘downstate’) characterized by lower firing rates (~1 spikes/s) and a more hyperpolarized membrane — likely due to a stronger inhibition from the dFSB (Donlea et al., 2018). In this downstate, we assumed that helicon cells’ resting membrane potential is closer to the chloride reversal potential, which makes them less responsive to inhibitory input, while still responsive to excitatory input. To implement helicon’s activity states in a current-based neuron model, we assumed a decrease of the inhibitory input from R5 to helicon at night (compared to the day) whereas excitatory input was unchanged **(Table 2)**.

To investigate coherent activity of R5 and helicon networks (**Figure S5-1, H and I**), we connected an initially desynchronized R5 network to helicon in the morning and night configurations and evaluated the synchronization within and between networks. In this case, R5 network was modelled with a low excitatory coupling to prevent synchronization when decoupled from helicon.

| Coupling | R5 – R5 | R5 – Helicon | Helicon – Helicon | Helicon – R5 |
|---|---|---|---|---|
| **Day**<br>$I_{exc}^{*}$<br>$I_{inh}^{*}$ | $0.051 \pm 0.017$<br>$-0.010 \pm 0.004$ | $0.830 \pm 0.102$<br>$-0.167 \pm 0.020$ | $0.0576 \pm 0.043$<br>$-$ | $0.423 \pm 0.079$<br>$-$ |
| **Night**<br>$I_{exc}^{*}$<br>$I_{inh}^{*}$ | $0.051 \pm 0.017$<br>$-0.010 \pm 0.004$ | $0.830 \pm 0.102$<br>$0$ | $0.0576 \pm 0.043$<br>$-$ | $0.423 \pm 0.079$<br>$-$ |
| $\tau_{exc}$ **(ms)**<br>$\tau_{inh}$ **(ms)** | 4<br>20 | 4<br>20 | 4<br>$-$ | 4<br>$-$ |
| **Synaptic connections** | $25.278 \pm 8.892$ | $103.762 \pm 12.704$ | $14.188 \pm 10.661$ | $52.938 \pm 9.877$ |

**Table 2.** Coupling between the neurons at day and night. Inhibitory and excitatory synaptic current amplitudes $I^{*}$ (mean $\pm$ standard deviation) and synaptic time constants. Synaptic currents are modelled as alpha functions. The numbers of synaptic connections (mean $\pm$ standard deviation) were estimated from the connectome (neuprint).

*1 Hz network stimulation*

In some simulations, a regular 1 Hz pulsed current was injected into all neurons of the R5 and/or helicon networks. The excitatory current pulses are modelled in the same way as the synaptic currents described in **Table 2** with $\tau_{exe} = 4$ ms. The current amplitudes were $I_{R5}^{*} = 3$ for the R5 network and $I_{Hel}^{*} = 15$ for the helicon network.

*Simulations*

For a single simulation, each neuron gets initially assigned at time *t*=0 a different, random membrane potential $v$ and recovery variable $u$. For $v(t = 0)$, random numbers are drawn from a range of -90 to -30 whereas for $u(t = 0)$ values are ranging from -11 to 0. The temporal resolution of simulations was 1 ms. The simulations for the PSDs were performed 10 times, and the mean and standard deviation were shown (**Fig. 5B, E**). In **Figure S5-1E**, the correlation coefficients of each value of $I_{exc}^{*}$ were computed 10 times, and the mean and the standard deviation were shown. In **Figure 5C**, the correlation coefficients between R5 and helicon for the nighttime and daytime simulation were computed for 10 simulations each. To compute the population activity or "firing rates" of R5 or helicon networks at each time step, we replaced each spike with a Gaussian (half width 300 ms) and averaged these filtered contributions of all neurons.

To determine possible compound oscillations of the networks, we computed the power spectral density (PSD). The power spectral density is computed by Welch's average periodogram method (matplotlib.pyplot.psd). We normalized the PSD for each network with respect to its number of neurons. To quantify the degree of synchronization of the activity of two signals (correlation coefficient), we computed their cross-covariance, which is a time-dependent Pearson correlation coefficient. We simulated network activity for 60 s and neglected the first 20 s in which the network activity stabilized. For quantification of synchronization within the R5 network, we again used filtered spike trains (Gaussian, 300 ms half width), computed the cross-covariance (at zero time delay) for all distinct pairs, and averaged across all pairs.
